# ModiDeC: a multi-RNA modification classifier for direct nanopore sequencing

**DOI:** 10.1101/2025.01.04.631307

**Authors:** Nicolò Alagna, Stefan Mündnich, Johannes Miedema, Stefan Pastore, Lioba Lehmann, Anna Wierczeiko, Johannes Friedrich, Lukas Walz, Marko Jörg, Tamer Butto, Kristina Friedland, Mark Helm, Susanne Gerber

## Abstract

RNA modifications play a crucial role in various cellular functions. Here, we present ModiDeC, a deep-learning-based classifier able to identify and distinguish multiple RNA modifications (*N*^6^-methyladenosine, inosine, pseudouridine, 2′-*O*-methylguanosine, and *N*^1^-methyladenosine) using direct RNA sequencing. Alongside ModiDeC, we provide an extensive database of *in vitro*-transcribed and synthetic sequences generated with both the new RNA004 chemistry and the old RNA002 kit. We show that RNA modifications can be accurately recognized and distinguished across different sequence motifs using synthetic data as well as in HEK293T cells and human blood samples. ModiDeC comes with a graphical user interface that allows easy customization and adaptation to specific research questions, such as learning and classifying additional RNA modifications and further sequence motifs. The reproducibility across samples, together with the low rate of false positives, underscores the potential of ModiDeC as a powerful tool for advancing the analysis of epitranscriptomes and RNA modification.

## Introduction

Chemical alterations of RNA molecules can affect their structure and stability as well as their interactions with proteins^1–4^, and stabilize or alter their metabolism at several stages of the cellular RNA life cycle^5^. Specifically, chemical modification of single nucleotides can interfere with RNA-dependent mechanisms involved in cellular localization, gene transcription, and RNA processing and localization^6^, all of which are crucial for protein production and the growth and development of eukaryotic cells^7–11^.

More than 170 different chemical modifications of RNA have been identified to date, giving us a glimpse of the complex epitranscriptome landscape^12^. Amongst the most prominent examples of RNA modifications are pseudouridine (Ψ)^2,13^, inosine (I)^14,15^, 2′-*O*-methylguanosine (Gm)^4,16^, and the N-methylation at position 1 and 6 in adenosine (m^1^A and m^6^A ^17,18^, respectively)^19^. For each of the before mentioned modifications it is known that an abnormal accumulation or an insufficient level of RNA modifications can cause severe cellular dysfunctions and diseases^25,26^. The spectrum of human pathologies triggered by RNA modification disorders ranges from developmental disorders and immunological deficits to neurodegenerative diseases and cancer in its various manifestations^8,22–25^.

Several methods have been developed for investigating the effects of RNA modifications on the transcriptome, including MeRIP-Seq^26^, m6ACE-Seq^27^, Pseudo-seq^28^, miCLIP^29^, and GLORI^30^. These methods combine antibodies or chemical treatments with next-generation sequencing for detecting and characterizing transcriptome-wide RNA modifications in short-reads. Despite the significant discoveries made using these methods, short-read sequencing technologies frequently struggle to accurately capture the diversity RNA modifications^1,31,32^.

Direct RNA sequencing (DRS) is currently revolutionizing the field of epitranscriptomics research as a technology that enables the identification of RNA modification at single-nucleotide resolution in long reads^33–39^. In nanopore sequencing, the passing of modified bases through the nanopores causes substantial changes in the expected current signal, enabling their identification, as impressively demonstrated in recent years^40–42^. In principle, DRS-based methods for detecting modifications can be categorized as comparative or *de novo* detection. Comparative methods such as xPore^43^ and DRUMMER^44^ compare negative controls with DRS analysis to detect RNA modifications with good performance. However, their requirement for unmodified control data limits their application. De novo methods such as m6Anet^45^, nanom6A^38^, DENA^46^, mAFiA^47^, or Penguin^48^ are based on training personalized deep neural networks using labeled datasets from sequences synthesized *in vitro* or RNAs transcribed *in vivo* to obtain ground-truth labels for modifications. These methods achieve single-base resolution for the analysis of a selected RNA modification, thereby demonstrating a new way to reliably predicting RNA modification. In this respect, direct RNA sequencing (DRS) holds great promise for opening up previously unknown areas of human disease diagnostics and biomarker development, and for bringing a technology-driven revolution to the field of epitranscriptomics.

However, despite the great progress, wider use of the technology has been hampered by several methodological obstacles, in particular the still significant inaccuracy of predictions in biological data, the lack of large-scale gold standard datasets for validation and the evolving sequencing chemistry. The latter is especially pertinent, as it renders the prediction tools that were specifically trained on certain features of one chemistry completely unsuitable. As a result, analyzing the epitranscriptomic landscape for different types of RNA modifications with DRS remains a challenge^49^. Several recent studies have, for example, highlighted the importance of being able to simultaneously detect multiple RNA modifications^50,51^ and the growing potential of direct RNA sequencing for investigating diverse RNA species and their modifications, including tRNA^52^, rRNA^53^, and other non-coding RNAs^54^. One example is a pioneering study focused on the interplay between m^6^A and Ψ in mRNA^55^. Using a specialized DRS pipeline, they were able to observe both the different effects of these two RNA modifications in total and polysome-associated mRNA and a change in translation efficiency related to the RNA modifications.

These distinctions between several modifications and the possibility to optimize the prediction-accuracy for specific regions or specific RNA types are precisely the technological advances needed if we are to fundamentally understand how a complex multitude of RNA modifications regulate health and disease. In particular, clinical RNA diagnostics requires reliable and easy-to-use options for the detection of RNA modifications that are integrated into standard software.

To overcome this challenge, we developed the Modification Detector Tool (ModiDeC). This flexible and customizable deep neural network can detect, classify and quantify different modifications. ModiDeC should be considered less as a competitor to other modification base callers that are designed to perform transcriptome-wide scans but more as a customizable tool that can be trained to be a highly specific classifier for certain regions, motifs, and modifications of interest. To offer the possibility of personalized training for individual research questions, we developed a GUI that allows the user to retrain ModiDeC to search for user-defined specific sequences or motifs without having to directly interact with the source code.

We demonstrate the performance of ModiDeC in analyzing synthetic sequences and selective RNA pseudouridylation mediated by synthetic designer organelles^56^ in HEK293T cell lines. In addition to synthetic approaches, ModiDeC was applied to ribosomal RNA data obtained from HEK293T cells and we verified its performance in detecting RNA modifications in human blood samples. Finally, we demonstrate the extendibility of ModiDeC by adding m^1^A to the pool of classifiable modifications. The exemplified fields of application in this study reveal the potential of ModiDeC as a personalizable high-accuracy multi-RNA modification classifier which can be extended and optimized as required for specific questions.

## Results

### Creating a training data set for ModiDec

To train a neural network to identify multiple modifications at different sequence motifs and distinguish them from one another, the neural network must be presented with various modifications in different sequence contexts. In this training dataset, the sequence, location, and modification type must be precisely known for each construct to form the so-called “ground truth”. To provide such training data, we designed and sequenced chemically synthesized oligonucleotides, allowing us to pick and choose the modified nucleotide as well as the sequence context. By ligating those oligonucleotides into a larger construct suitable for direct RNA sequencing, which performs optimally with long reads, we created multiple point-modified sequences with complete information regarding their modification status. Details on the design of the training data are provided in Section SI 1 of the Supplementary Information. Ten oligonucleotides were created per modification type (m^6^A, Gm, I, and Ψ; see Supplementary Table 1) to give a total of 40 different oligonucleotides. Each oligonucleotide contained a 9-mer motif flanking the modified nucleoside, along with the same unmodified 9-mer sequence, such that the training included both modified and unmodified motifs.

For oligonucleotides containing m^6^A, the sequences were based on the evolutionarily conserved DRACH motif^17,57^. To ensure that ModiDeC distinguishes modifications based on their specific signal rather than the surrounding sequence context, one DRACH-motif-containing sequence was selected to represent a single oligonucleotide motif across all modification types. For the remaining motifs, we either mirrored known modification sites in the human transcriptome, as reported in the Modomics database^14^, or chose naturally occurring sequences but without a known modification background. A list of all sequence motifs is provided in Supplementary Table 1. The oligonucleotides were then incorporated between two *in vitro*-transcribed RNA sequences via splinted ligation, creating a larger sequence construct ready for sequencing (see Methods and Supplementary Fig. 1b). Following this strategy, for all 40 modified constructs we also created a corresponding negative control without any modification.

The RNA004 flow cell was introduced by Oxford Nanopore Technologies for use on their DRS platform in 2023. Its new chemistry differs significantly from the SQK-RNA002 kit in several ways^58^, notably in *k*-mer length, dwell time, current level, and translocation speed. Because ModiDec is intended for use with data acquired using the old RNA002 or the new RNA004 sequencing chemistry, we double-sequenced all 40 constructs (once with the SQK-RNA004 kit and once with SQK-RNA002), together with the unmodified control, thus creating two identical data sets that differed only with respect to the kits/flow cells used in DRS. This allowed us to train an accurate model for each chemistry, giving the user the flexibility to analyze any DRS data sets with ModiDeC, regardless of the chemistry used (Fig. 1a).

**Fig. 1:**
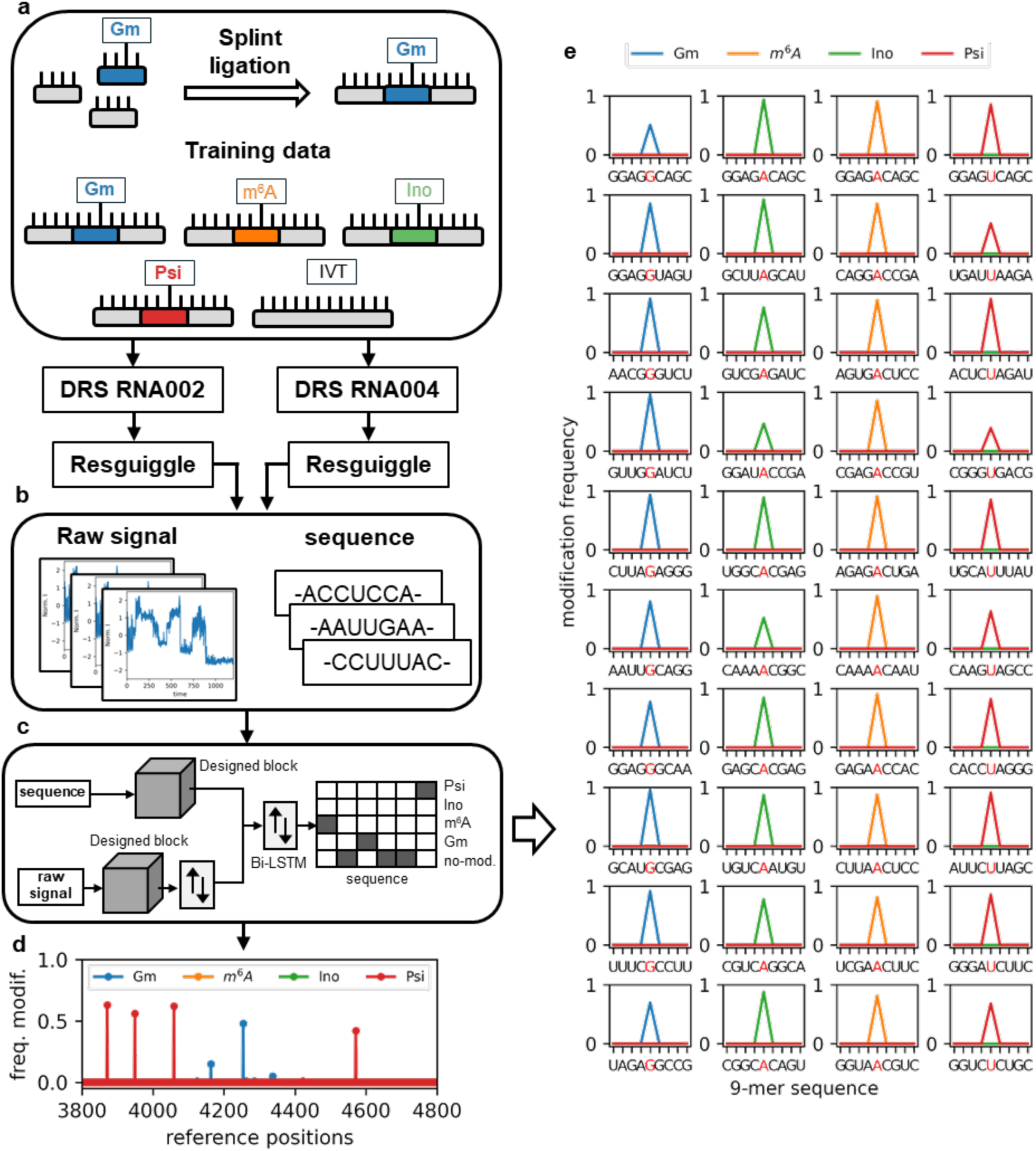
Schematic representation of the workflow - from data generation to pre-processing, model training and prediction. a) Representation of oligos used for training ModiDeC. Modified and unmodified labeled reads were used for the training. b) After base calling, data are re-squiggled to the reference and chunks with respective references are sent to the neural network. c) Scheme of the ModiDeC neural network. d) Example output after neural network analysis. e) The modification frequency on the test data for the 40 motifs used in this work (RNA004 kit). The red letter is the modified base. Each panel shows the predicted frequency of modification (y axis) with the respective modification type and modified motif reference (x axis) in the oligos. For clarity, we visualize only the core motif of the 9-mer. The red letter indicates the modified base.

In addition, we designed five constructs for the initial testing of ModiDec. These constructs contained either no modification, one in a previously unknown motif, or two distinct modifications in motifs the model had been exposed to in training. These sequences were also generated by splinted ligation, and in the case of doubly modified constructs, the final construct sequence alternated between the oligonucleotides and the IVT sequences to ensure sufficient distance between the modified oligos. For details of the ligation process, see Methods and Supplementary Fig. 1c.

### Training and validation of ModiDeC neural network

ModiDeC is a personalized two-input neural network that combines long short-term memory layers (LSTM) with a newly designed Inception-ResNet block to classify multiple RNA modifications simultaneously on the data. In detail, the neural network re-maps the sequence by analyzing the signal, and each sequenced nucleotide is classified as either “unmodified”, Ψ, Gm, I, or m^6^A. ModiDeC was trained using the 40 splint-ligated sequences, each containing one modified oligonucleotide (Supplementary Table 1 and Supplementary Fig. 1a,b). Thirty thousand reads per modification and motif were used for training. This number was chosen following a pre-analysis investigating the change in prediction accuracy as a function of altering the amount of training data. As expected, the percentage of correctly accessed modification sites increases with the amount of training data (see Section SI 2 of the Supplementary Information for more details). However, 90% accuracy can already be achieved with approximately 5,000 reads. With more reads, the accuracy increases only minimally and almost reaches a plateau at 30,000 reads.

After base calling, each read was re-squiggled to the reference, cut into chunks of 400 time points, linked to their sequence, and then used for the input of the neural network (Fig. 1b, see also Methods for more details). For each chunk, an output label for the neural network was created in parallel. The output of the neural network is a one-hot encoding representation, where each base is classified between “unmodified” or one of the four modification types (m^6^A, Gm, I, and Ψ). In total, more than 40 million labeled chunks were used to train ModiDeC (Fig. 1c). The final accuracy of the neural network to identify the correct modification position after the training phase was higher than 91%. An example of a ModiDec analysis output is shown in Fig. 1d. After selecting a pod5 file, the corresponding aligned bam file, the reference, and the number of reads to analyze, ModiDeC re-maps each read and finally reports the frequency of predicted modifications at the single-nucleotide level for each position in the reference.

We validated ModiDeC on a new dataset that contained all the motifs and modifications used during training but had never been seen or analyzed by the neural network. Five thousand reads were used for each oligo. All test data were sequenced with SQK-RNA004. Fig. 1e shows the results of the validation analysis. The network correctly identified the position and type of modification 100% of the time for the 40 sequence modification combinations. The rate of correct quantification of the occurring modification (detection of the modification frequency) was 81%. However, this average prediction accuracy is both motif- and modification-dependent. Although we can report impressive prediction accuracies of almost 100% for some motif–modification combinations, others were obviously more difficult to detect. This might also be partly due to the sequencing runs differing slightly in quality. In each individual case, however, a modification was clearly identified (even if correct quantification was not always achieved). A detailed overview of true-positive, false-negative and modification miscalling predictions for each motif modification combination is reported in Supplementary Table 2).

Particularly interesting in this context are the results for the DRACH motif AGACA in which a different modification was introduced at the same position in each case (Fig. 1e, first row). The data show that ModiDec correctly predicts that the A base in this DRACH motif is modified. However, because we have inserted different modifications at the same motif, we can exclude that ModiDec has learned only from the sequence in the training data that the DRACH motif, for example, stands for m^6^A modifications. Instead, we can show that ModiDec can correctly distinguish and classify between several modifications, even if a new modification appears on an actual “typical” motif for another modification. Because ModiDec makes an independent prediction for each individual read at the single-molecule level, even a mixed prediction could also have been made. However, ModiDec was able to distinguish precisely between the four modifications in every case. To determine the false-positive prediction rate precisely, we validated ModiDec on the negative control data. We observed that ModiDeC has an extremely low false-positive prediction rate between 0% and 1% (Supplementary Fig. 3).

Next, we wanted to test if similar results could be reproduced in the dataset sequenced using SQK-RNA002. To this end, we retrained and validated the neural network using the same datasets but sequenced with the old chemistry. Also, with SQK-RNA002-acquired data, ModiDeC shows great ability to predict precisely the position and the corresponding modification for each individual motif–modification combination (for details of this analysis, see Supplementary Figs. 4 and 5 and Supplementary Table 3). Interestingly, the SQK-RNA002 dataset also shows that some of the combinations can be excellently predicted, but for others, the network tends to more frequently misclassify modifications as “unmodified”. However, these are not the same combinations as in SQK-RNA004, which again suggests a difference in the quality of the sequencing performance rather than a systematic bias in the network’s predictions. Regarding the determination of the false-positive prediction rate, a similar result emerges from an analysis of the unmodified control data sequenced with SQK-RNA002 to that we observed with SQK-RNA004. Although we can discern a slight difference between the performances of the models trained with the old and new chemistries, the numbers of false-positive predictions are still low, ranging between 0% and 5.5%.

In conclusion, the new RNA004 chemistry gives a slight improvement in detection accuracy, but ModiDec works reliably with both the old and new chemistries. Of articular note here is the high accuracy of the position prediction (base-level resolution), as well as almost perfect classification. Since no major differences were found between the two kits in terms of ModiDec’s performance, and the SQK-RNA002 kit is no longer manufactured, we decided to focus on the results of the SQK-RNA004 kit for further analysis in this study.

Next, we tested ModiDeC on five RNA constructs previously unknown to the network and with varying complexity of modification. These constructs either carried no modifications at all (Fig. 2a), carried one modification (Fig. 2b,c), or contained two adjacent modifications at a specific position in the sequence (Fig. 2d,e). These five constructs were designed as a random sequence in which a total of six of the motifs examined here (unmodified or modified) were inserted into the sequences at specific positions (Fig. 2). For the unmodified reads, the neural network correctly returned zero modified positions (Fig. 2a). For the singly modified reads, ModiDeC identified constructs composed of two oligos, one of which contained a position (73) modified as Ψ (Fig. 2b,c) Oligos used for the construction were identical in sequence between constructs and therefore only differed by which oligonucleotide contained the modified position. ModiDeC was able to identify correctly the positions modified and classify perfectly the type of modification linked to those positions with an accuracy higher than 86%. The fact that both constructs shared identical oligonucleotide sequences highlights that ModiDec accurately detects the state of modification rather than assigns them based on sequence patterns. Next, the systems of greatest complexity—those doubly modified—were analyzed using ModiDeC (Fig. 2d,e). For both oligos, the neural network was again able to identify correctly the positions of modification and classify the modification type linked to those positions with high accuracy (> 85% of all modifications could be correctly detected on a single-nucleotide level).

**Fig. 2:**
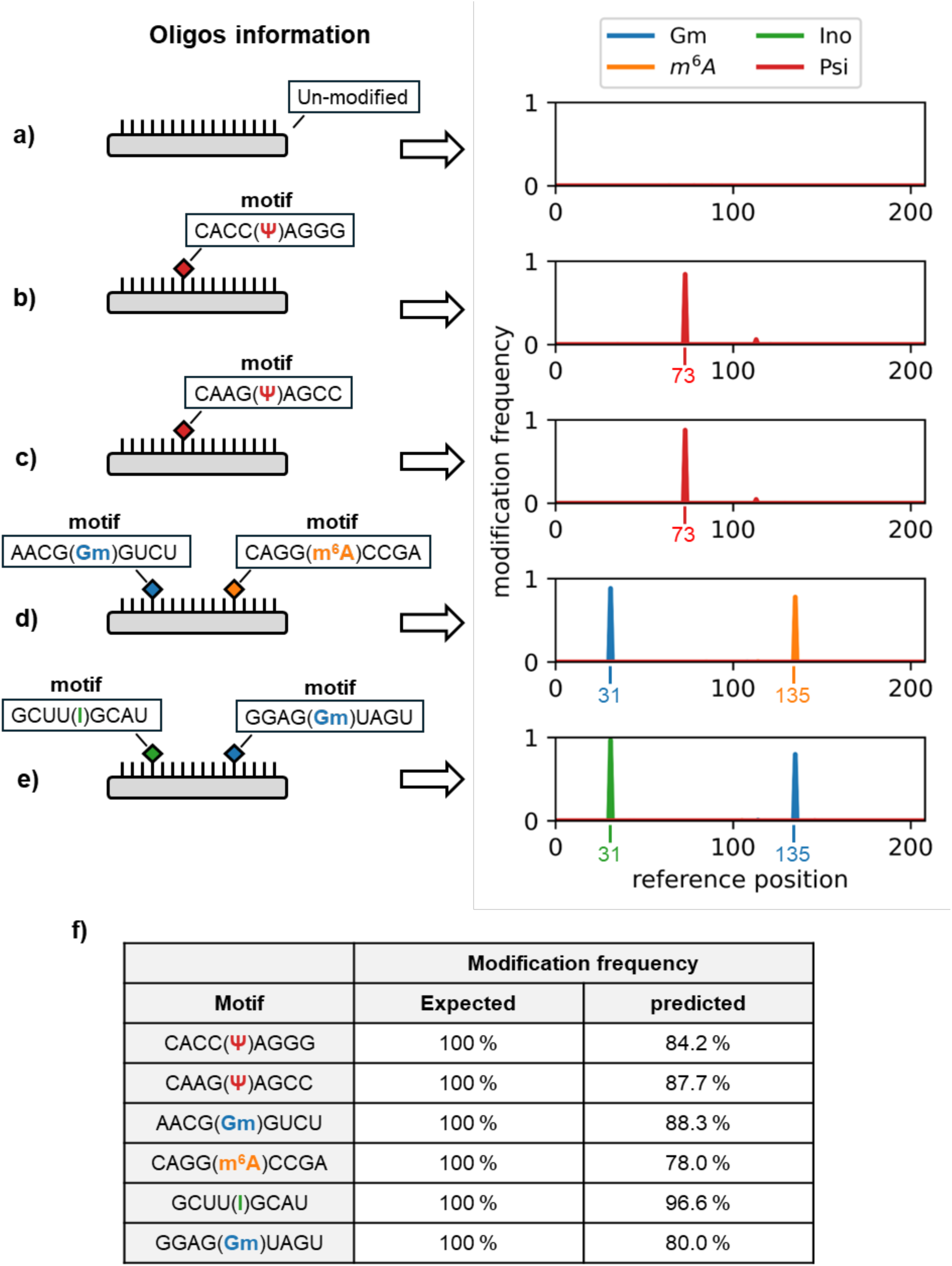
ModiDeC analysis on new RNA oligos for reads with varying complexity. (a) unmodified, (b,c) singly modified and (d,e) doubly modified reads. (f) Table showing the expected and ModiDeC-predicted modification frequency for each motif.

From the test analysis we observed that ModiDeC always detected the modification position with a high frequency of modification. However, it underestimated the expected value by about 10% on average.

### HEK293T cell analysis: selective organelle and ribosomal RNA

To test the ability of ModiDeC to detect and quantify modifications in physiological data sets, we continued our study using data obtained from HEK293T cells.

The first example is an already characterized “optimized RNA-editing organelles (OREO)” system⁵⁶, which in combination with guide RNAs can selectively pseudouridylate a target RNA. The OREO system has been tested on a selectivity reporter harboring two target positions in mCherry and EGFP sequences, in which the former is preferentially modified based on an optimized recruiting system of the organelle. The selectivity of pseudouridylation was confirmed by measuring the amount of pseudouridine in specific target sites of mCherry (position 565) and EGFP (position 115) mRNAs in HEK293T samples transfected either with the designer organelle (sample O) or without the organelle (sample nO) using DRS together with the pseudouridine detection model (Oxford Nanopore Technologies).

ModiDeC was used to reanalyze the EGFP and mCherry data sets to track and quantify pseudouridine in three of the published samples, namely the sample with the organelle (O), without the organelle (nO), and a control sample without any targeting system (C) for both RNA target sites in the selectivity reporter. For this purpose, ModiDeC was previously exposed to the pseudouridine target positions—modified both in training and testing phases (see Figs. 1e and 2b,c). The results are shown in Fig. 3a. We found that ModiDeC correctly predicted the expected position (red numbers in Fig. 3a) and assigned a Ψ modification to the U nucleotide. Comparing the nO sample with the O sample, we observed a reduction in Ψ from 36% to 13.8% (2.61-fold) at the EGFP site, whereas for mCherry the reduction was from 73% to 56% (1.3-fold). The control sample also showed Ψ peaks for EGFP (∼11%) and mCherry (∼3%). These values are consistent with those published and experimentally validated in the original work by Schartel et al. in 2024^56^.

**Fig. 3:**
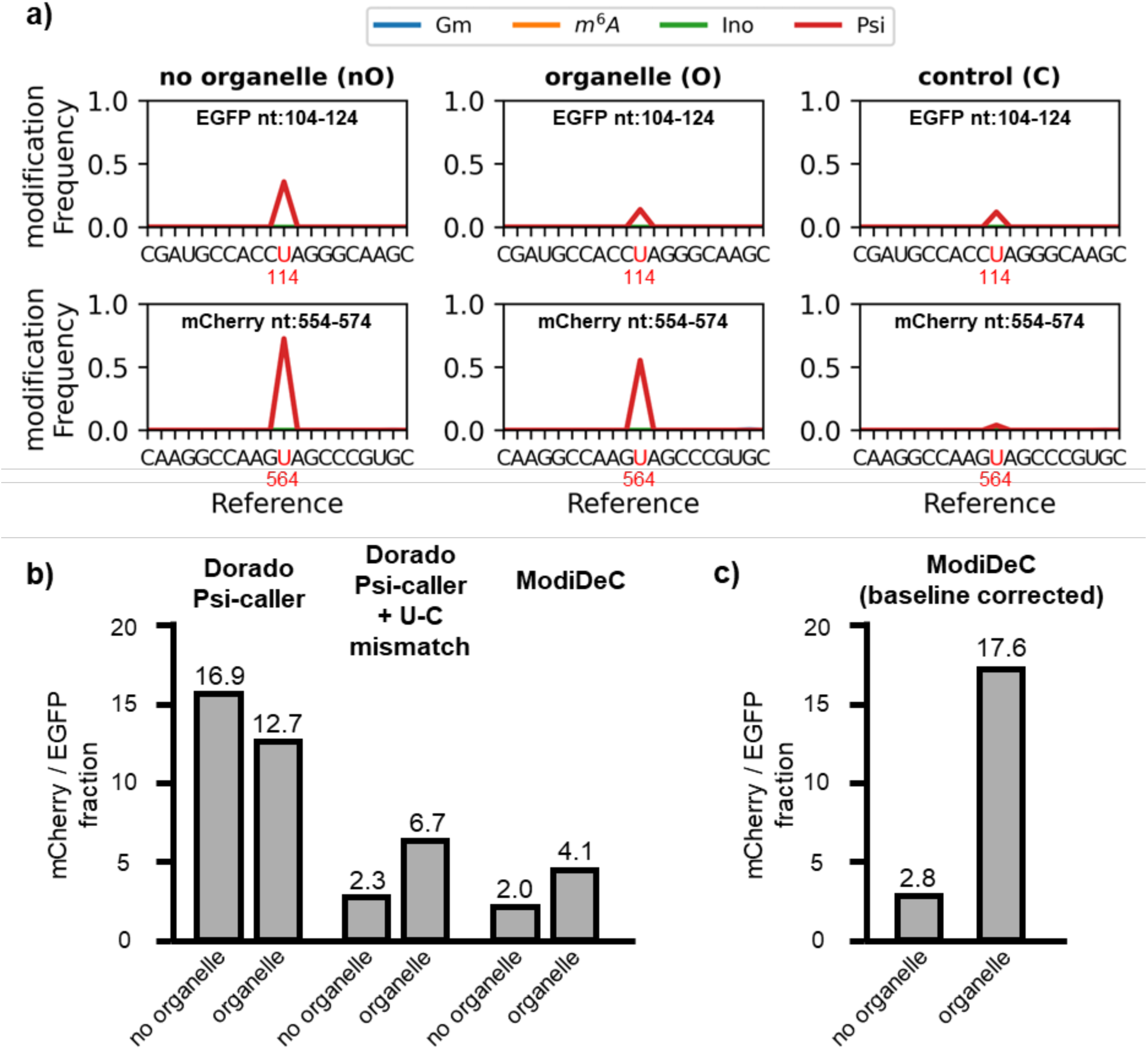
HEK293T OREO system analysis using ModiDeC and Dorado. a) ModiDeC analysis on HEK293T RNA samples without organelle (left), with organelle (center) and control (right) for EGFP (top row) and mCherry (bottom row). Red numbers and letters indicate the expected modified base and position. b) mCherry/EGFP ratio calculated for organelle and no-organelle systems using Dorado pseudouridine model (Dorado-*Ψ* -caller), Dorado-*Ψ* -caller + U–C mismatch, and ModiDeC. Expected trends can be tracked with Dorado-*Ψ* -caller + U–C mismatch and ModiDeC. c) mCherry/EGFP ratio from ModiDeC after subtracting the control (baseline correction) for organelle and no-organelle systems.

Next, we calculated the mCherry/EGFP pseudouridine ratio, a measure of organelle selectivity, for both OREO-containing and OREO-free systems for Ψ counts predicted by ModiDec and compared it to the ratio derived from the Ψ-detection model implemented in Oxford Nanopore Technologies’ most recent base-caller Dorado (Fig. 3b). Although that model was able to detect the Ψ sites, it did not reproduce the expected trend. By contrast, the predicted mCherry/EGFP pseuduridylation ratio was lower for the OREO-containing system (12.7-fold) compared to the OREO-free sample (16.9-fold; see Supplementary Fig. 6 for Dorado Ψ -caller values). This unexpected result can be mainly explained by the fact that the Ψ model from Dorado exclusively considers U-called sites and therefore overlooks mismatched bases. As shown by Schartel et al. and others, Ψ can cause the motif-specific base-calling error of a U-C mismatch in DRS data^41,56^. Only by manually counting the U-C mismatches and adding these values to the results of the Dorado pseudouridine caller, the expected increase in the mCherry/EGFP ratio from 2.3-fold to 6.7-fold was observed upon switching from the OREO-free to the OREO-containing system (see Supplementary Fig. 6 for Ψ and U-C mismatch Dorado values).

By contrast, ModiDeC was able to reproduce the increase in mCherry/EGFP pseudouridylation ratio in sample O versus sample nO, by considering the detection of Ψ only. We also calculated the mCherry/EGFP ratio from a ModiDeC analysis performed in the same way as in the original study^56^, where the Ψ values obtained from the non-targeting control system were subtracted (baseline correction) from the pseudouridine counts of both organelle and no-organelle systems (Fig. 3c). The baseline-corrected ModiDeC analysis showed fold-changes of 2.8 and 17.6 for no-organelle and organelle systems, respectively, which is consistent with the validating bisulfite-induced deletion sequencing (BID-seq) values reported^56^.

Next, we analyzed a fragment of 18S ribosomal RNA (Fig. 4a; for a complete structure of 18S, see Supplementary Fig. 7). We chose this region because it contains documented modification sites^59^. In our synthetic training dataset (Fig. 1d) we integrated five motifs for Ψ and Gm covering known and isolated modification sites on 18S rRNA (see table in Fig. 4b). We analyzed two *in vitro* samples (sample 1, rRNA from the cytoplasm; sample 2, rRNA from the nucleus) of the 18S derived from HEK293 cells together with synthetic 18S IVT reads as a negative control (Fig. 4c). Because both samples originate from the same cell line, albeit from different compartments, we expect a conservation of modification sites at varying frequency of modification. The ModiDeC analysis of samples 1 and 2 shows a highly congruent prediction of Ψ- and Gm-modified sites, precisely matching the trained motifs (Fig. 4c, upper and middle panels). The prediction of modification frequency showed the expected variations between comparable loci of different conditions (e.g., peaks 3949 and 4571; see Fig. 4b), while maintaining similar dynamics between distinct loci within a sample. The classification of Ψ- and Gm-modified sites was confirmed for every predicted site. In addition, the analysis of 18S IVT data (Fig. 4c, bottom graph) shows a drastic reduction of the modification frequency at modification sites matching trained motifs, indicating a low false-positive rate.

**Fig. 4:**
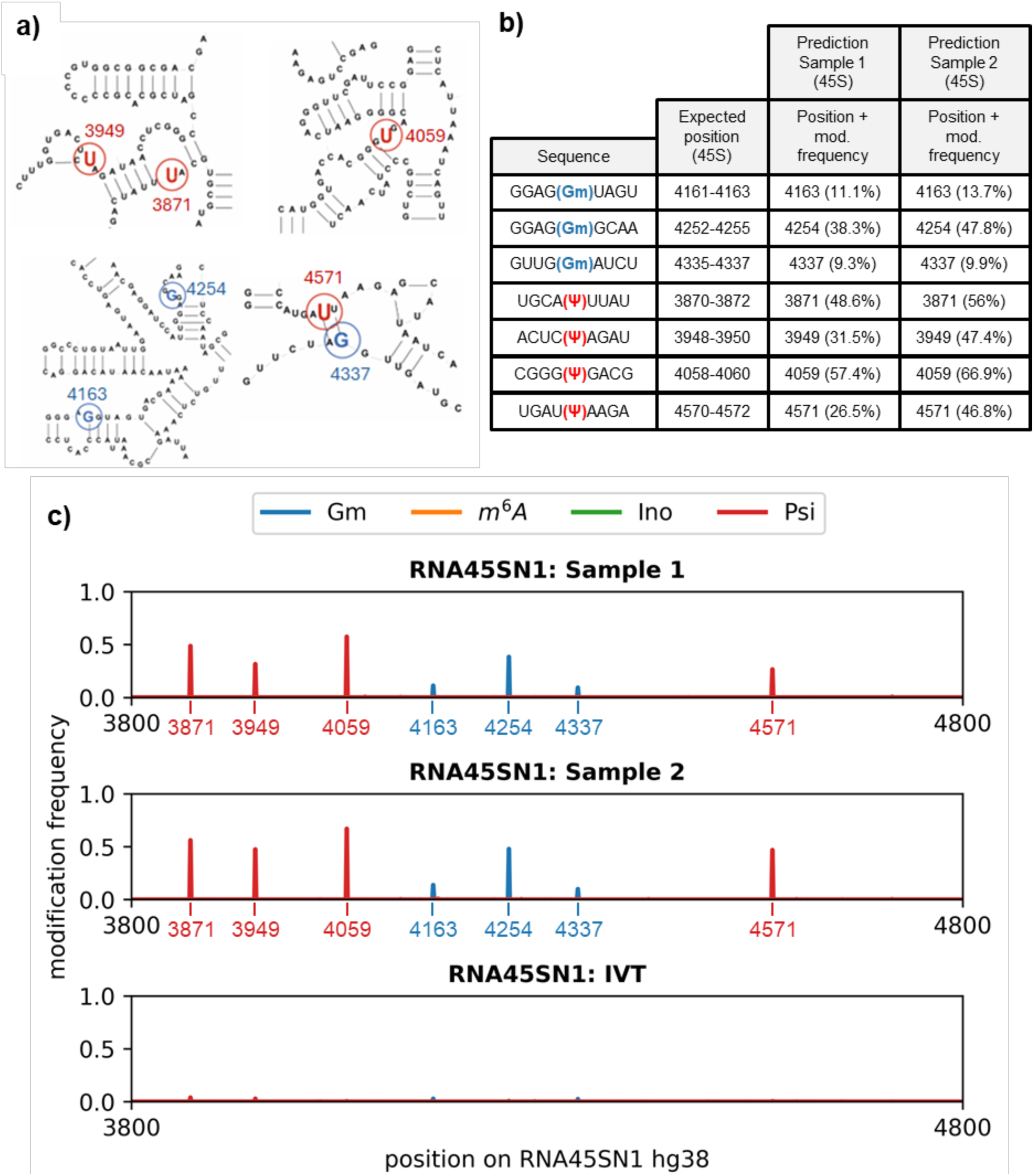
Analysis of 18S ribosomal RNA. a) RNA45SN1 RNA structure. The highlighted letters are the modified bases recognized by ModiDeC (red = *Ψ*, blue = Gm). The numbers indicate the reference position. b) Table showing expected motif sequence, expected position (45 S) and the ModiDeC analysis on samples 1 and 2. ModiDeC correctly predicts the positions, showing significant values for modification frequencies. c) ModiDeC predictions for RNA34SN1 hg38 for sample 1 (top), sample 2 (middle), and IVT (bottom). Both samples 1 and 2 show distinct *Ψ* and Gm peaks compared to IVT.

In summary, the analysis of the physiological HEK293T cell data in the two examples above validates the performance and customizability of ModiDeC. Once trained on a specific context, it can be used for the quantification of modifications in biological systems (Fig. 3) as well as for precise detection and classification of position (Fig. 4). ModiDeC also showed impressive reproducibility in its results.

### Detecting RNA modification in human blood

Next, we were interested in the general applicability of ModiDeC for diagnostic purposes. To this end, we tested ModiDeC on human transcriptome data from the most common sample type in routine diagnostics, that is, peripheral blood. It is important to mention that ModiDeC should be considered less as a competitor to other modification base callers that are designed to perform transcriptome-wide scans but more as a customizable tool that can be trained to be a highly specific classifier for certain regions, motifs, and modifications of interest.

The following strategy was chosen to assess the accuracy of the forecasts: We analyzed and compared both human peripheral blood from a healthy subject and a generated whole-transcriptome-wide IVT data from the same individual as a negative control. ModiDeC predictions were then validated with available experimental data from the literature.

As an example, we chose the well-studied and annotated TP53 tumor suppressor, responsible for multifunctional transcription factors, such as DNA repair or cell cycle arrest, and known to interact with the epitranscriptomic network of m^6^A^60–62^. ModiDeC was used to analyze the 33 transcripts linked to the TP53 gene with the purpose of identifying m^6^A RNA modification sites that are indexed in the RMBase v3.0 database^63^. It should be noted that the annotations of the RMBase v3.0 have not been experimentally validated for peripheral blood, but only for human cell lines (HEK293). Although, a tissue-specific methylation landscape must be assumed, we have nevertheless used this resource for validation. Since no database for m6A sites in human peripheral blood currently exists, this resource is the one that most closely matches our data in the literature.

The database reports 49 possible m^6^A RNA positions on those transcripts linked to TP53. Aligning these positions with ModiDeC m^6^A training motifs, the analysis could downstream to 12 reported m^6^A RNA positions that have a strong overlap with the training motifs which we have therefore considered in the following.

These 12 possible m^6^A sites were correctly detected by ModiDeC, and m^6^A peaks were predicted at four of these sites in three different transcripts (Fig. 5a-e). The dashed black lines in Fig. 5b-e indicate the expected position of the m^6^A site as reported in the RMBase v3.0 database. All four positions have an exact overlap with the expected position. Moreover, the negative controls rule out a false positive at the same positions in the IVT data. In addition to the four annotated m^6^A sites, ModiDeC has detected three further m^6^A positions (Supplementary fig. 8) that might be very interesting candidates for further experimental validations.

**Fig. 5:**
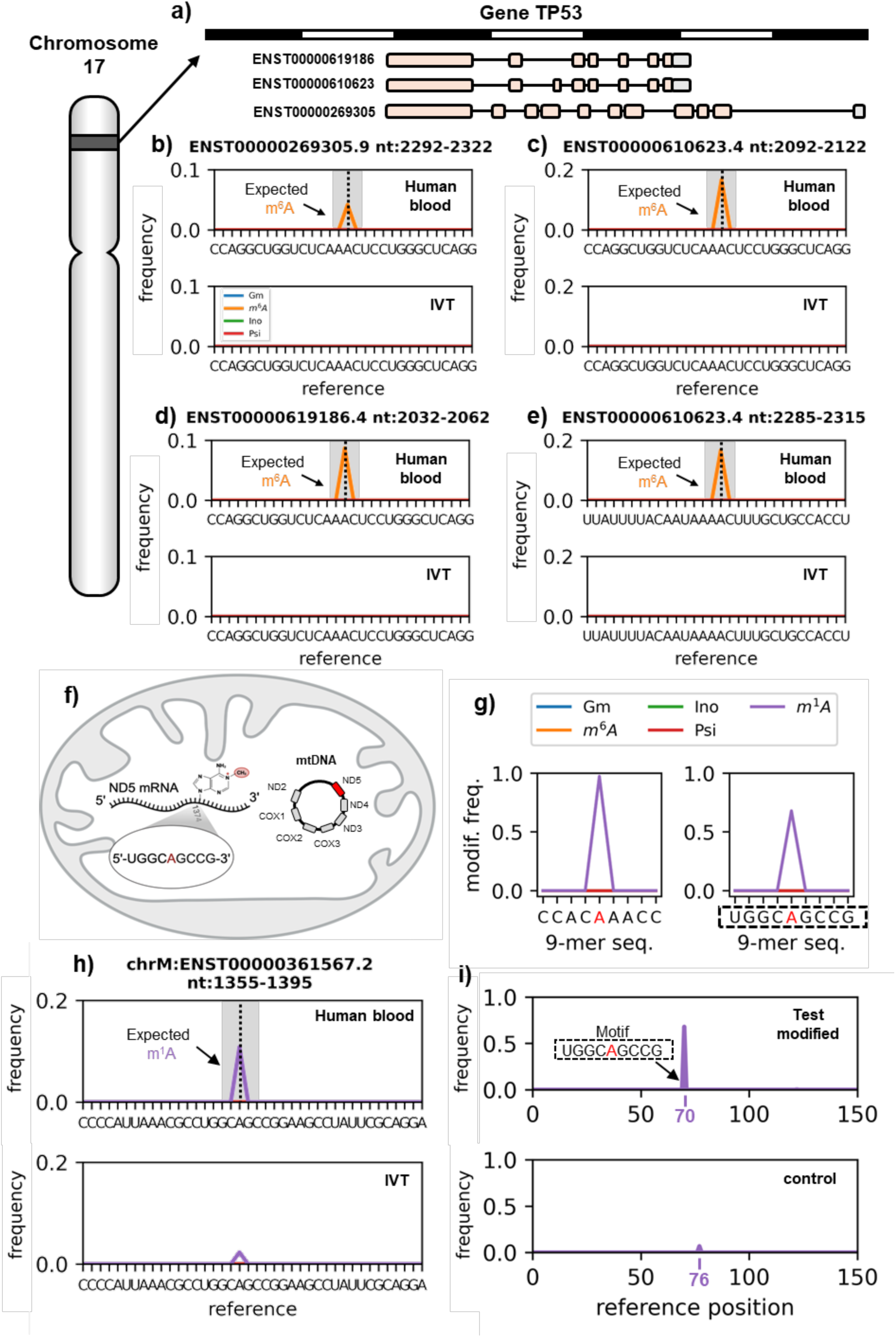
Detection of site-specific RNA modification in human peripheral blood samples. a) TP53 gene cartoon representation from chromosome 17. b-e) ModiDeC analysis on four different transcripts linked to the TP53 gene for the human blood sample and IVT data sets. While modification peaks are detected at the expected positions, w predictions on the IVT are a baseline. *f*) Mitochondrial ND5 RNA showing the motif containing the m^1^A modification (red “A”). *g*) ModiDeC validation on two different 9-mers (9-mers are shown in the x axis). The red letter indicates the modified base in the validation data. *h*) ModiDeC results on modified test data *(top) and* control sample *(bottom).* The peak at position 70 indicates a modification classified as m^1^A. The sequence is the mitochondrial RNA motif, with the modified base indicated in red. *i*) ModiDeC analysis on a mitochondrial human blood sample *(top) and* mitochondrial IVT data *(bottom)*. ModiDeC can detect the expected m^1^A peak associated with the mitochondrial ND5 RNA.

To show that ModiDeC is also suitable for predicting the existence of all four modifications (m^6^A, Gm, Ψ and I) simultaneously in our human blood sample, we also analyzed a second group of transcripts, together with the matching IVT data. (See Supplementary Figure 9 and 10). Here again, we were particularly interested in transcripts that are potentially clinically relevant. For example, the transcripts shown in Supplementary Figure 9a-e are derived from genes known to be dysregulated in cancer. For example, CD147 (Supplementary Fig. 9a) plays an important role in the development of various malignant tumors^65^, while RAB11B (Supplementary Fig. 9e) is a member of the RAS oncogene family^64^. Regarding potential validation, we were able to validated the m^6^A sites with the help of the m^6^A-Atlas in the due course^64^. According to the table in Supplementary Fig. 9f, all the predicted m^6^A motifs were confirmed by the reference database; most of the predicted positions match the annotated ones exactly. Only two of the predicted positions are less precise, having positional shifts of 1 and 6 nt. However, we cannot confirm whether this observed deviation is due to ModiDeC making an inaccurate shift in the prediction or whether the annotation is imprecise at this point due to inexact experimental resolution.

To the best of our knowledge, there are currently no data resources for the other predictions that we could have used to reliably confirm the validations for inosine and GM. Therefore, we refer to these only in the supplement and note that, from our point of view, they are highly interesting points for further targeted experimental validation steps. Following this strategy, we can therefore envision ModiDeC as a tool for the targeted prediction of previously unexplored modifications.

Another interesting finding is that ModiDeC obviously can also identify positions where the sequence differs slightly from the trained motif (Supplementary Table. 4). All listed positions contain at least one nucleotide that differs from the initially trained motifs (as listed in the Supplementary Information SI1). Some positions even contain partial combinations of the training sequences. This suggests that ModiDeC recognizes features of modifications in sequences that are even beyond the set of motifs used for training.

Next, we wanted to show that ModiDeC can also recognize further modifications, such as m^1^A sites. For validating the predictions in the biological data, we selected a specific N^1^-methylation of adenosine (m^1^A) on the mitochondrial *ND5 RNA*, which is described in the literature for its biomedical relevance^21^. Due to the methodological difficulties in using commercially available nonspecific antibodies against m^1^A, the sites of m^1^A RNA methylation predicted to date are highly controversial. Using a more stringent transcriptome-wide mapping approach, at single-base resolution, m^1^A was only detected in a low number of cytosolic mRNAs^19,65^. Irrespective of the approach used, numerous studies have clearly identified mitochondrial ND5 RNA modified with m^1^A at position 1374^19^. However, the current mapping approach is rather complicated and time consuming. The ability to identify m^1^A sites based on direct RNA sequencing would be ground-breaking for the field.

Therefore, we expanded the pool of classifiable RNA modifications by introducing two additional motifs into the ModiDeC analysis: the first was a randomly chosen natural motif without a known modification context (CCACm^1^AAACC), while the second was the specific motif of the mitochondrial ND5 RNA transcript sequence (UGGCm^1^AGCCG)^21^ (Fig. 5f). The full sequences, and that of the unmodified control, are shown in Supplementary Table 1. The predictions on the validation data (generated using the same splint–ligation strategy as before), together with the frequency of detection of these motifs in the evaluation data, are shown in Fig. 5g. For our modified validation dataset, the average frequency recognition was approximately 82% and the modified site was accurately detected in both cases. We also detected the m^1^A site in a new sample (having a sequence previously unknown to the network) that contains the motif with an m^1^A modification at position 70 (Fig. 5h, top) and compared it with the results from the unmodified control sample (Fig. 5h, bottom). ModiDeC correctly detected the position of m^1^A with a frequency of around 71% (Fig. 5h, top). In the control sample (Fig. 5h, bottom), almost every nucleotide is correctly classified as “unmodified” except a small peak with a frequency below 5% at position 76, which might indicate a threshold for the reliability of results from an m^1^A analysis.

After the successful training with the modified oligos, the m^1^A-expanded version of ModiDeC was then used to analyze the mitochondrial RNA that served as a template for the training motif. For this purpose, we again used the blood of the healthy subject and specifically targeted the mitochondrial ND5 RNA transcript (fig. 5i). ModiDeC analysis showed an m^1^A peak with a modification frequency around 11% at the expected position of 1374. The IVT data of the same transcript was analyzed, and an m^1^A prediction of less than 2% was found in the same position; we considered this a false-positive.

The generation of false positives by ModiDeC was also investigated by analyzing at random 160 IVT transcription sites (each with more than 80 reads per transcript) from the chromosomes here studied. We found that ModiDeC in 97% of cases has a specific false-positive region between 0% and 5%, while in 86% of cases a value between 0% and 3%. This rate is consistent with other analyses in this study. Although Based on these values, we suggest using ModiDeC for biological data sets with a threshold of 5% to almost wholly exclude false-positive peaks.

In Summary, on the human data ModiDeC was able to make precise and reproducible predictions for m^1^A and m^6^A for mitochondrial and TP53 (chromosome 17) transcripts respectively. Additionally, using synthetic oligos and human blood data, we demonstrated that ModiDeC can be easily extended with further modification classes (m^1^A) and motifs in addition to the initial 40 sequences on which it was originally pre-trained (see Supplementary Table 1).

### ModiDeC GUI: a graphical user interface for retraining the model

To enable users, regardless of their expertise in bioinformatics, to use and expand ModiDeC, we created a user-friendly pipeline that includes three graphical user interfaces (GUIs; Fig. 6). These GUIs facilitate the use of the model for analyzing specific locations or RNA-types to personalize it, or to retrain it without having to directly interact with the source code. The pipeline consists of three parts: “ModiDeC data curation”, “ModiDeC training”, and “ModiDeC analysis”. “ModiDeC data curation” (Fig. 6, top), supports the user in generating a data set for re-training the neural network. This GUI is highly customizable and allows the user to preprocess data by linking output labels to signal information for the proper generation of training data. The “ModiDeC training” GUI (Fig. 6h, middle) allows the selection of neural network parameters such as batch size and epochs. The “ModiDeC analysis” GUI can be used to visualize modification sites and corresponding modification frequencies on the reference sequence (Fig. 6, bottom). A more detailed explanation of the GUI inputs can be found in the Supplementary Information (Section 8). Additionally, a tutorial for using the three GUIs can be found on the GitHub page (see section “Code Availability”).

**Fig. 6:**
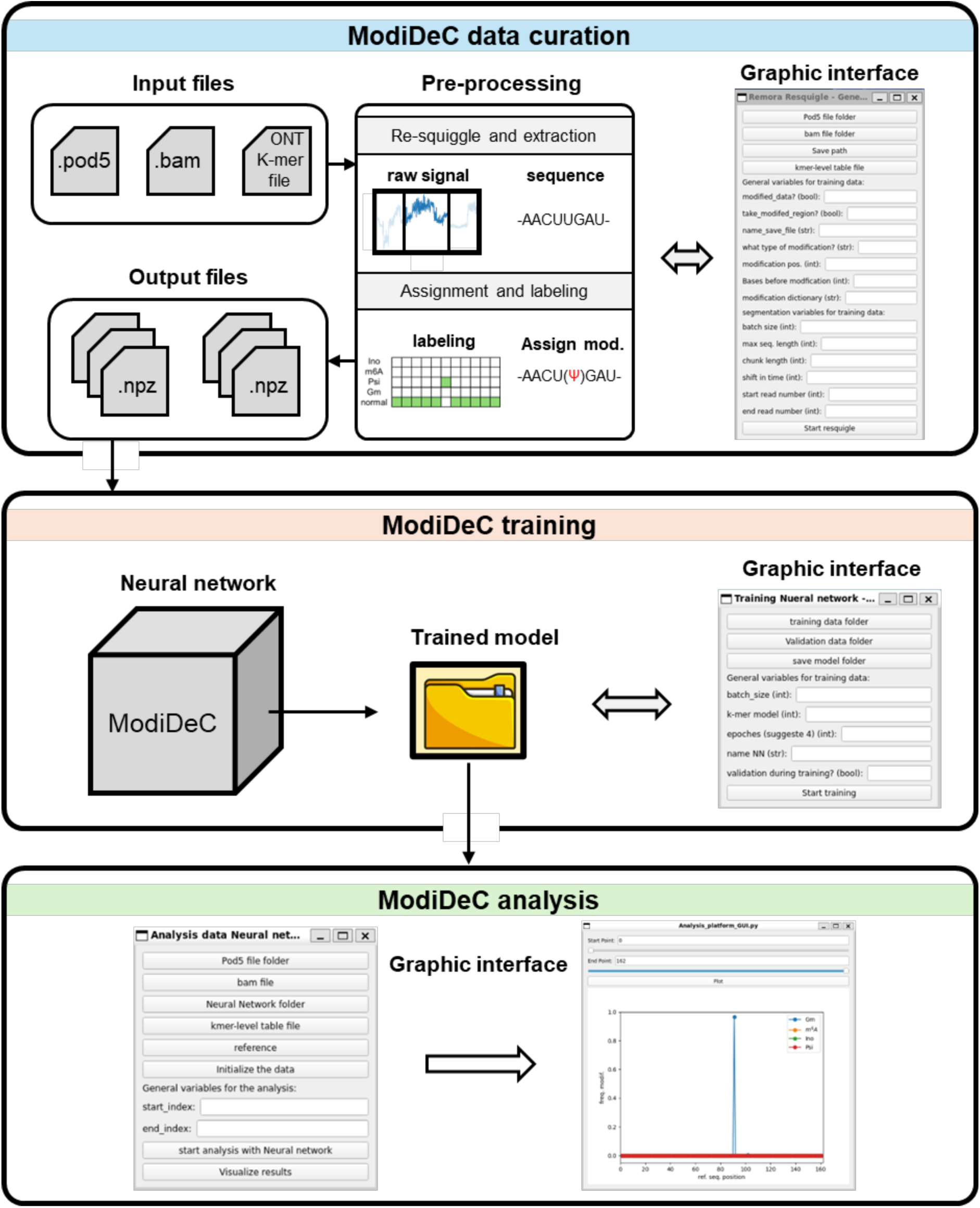
ModiDeC workflow to adapt the neural network to specific biological analysis using ModiDeC’s graphical user interface (GUI). Schematic representation of the ModiDeC’s GUI pipeline. The GUI is divided into three communicative sub-parts, created for retraining the neural network or directly using it for data analysis. Top) “ModiDeC data curation”. Raw data can be pre-processed to correctly create the input and output for the neural network training. Middle) “ModiDeC training”. Helps the user to easily retrain the ModiDeC for their problem. Bottom) “ModiDeC analysis”. Interface created to visualize and store the model analysis.

## Discussion

The neural network ModiDeC was developed to recognize whether an RNA nucleotide is modified at a specific position and assign it a specific modification. ModiDeC can be used with DRS data sequenced with an SQK-RNA002 sequencing kit, as well as with those acquired with the newer RNA 004 chemistry.

To demonstrate its performance, ModiDeC was initially trained on four different types of RNA modifications (m^6^A, Gm, I, and Ψ) with 40 different motifs (10 for each type of modification). Expanding this fundament, we demonstrated that ModiDeC can be simply trained for further motifs or new modifications, exemplified for a mitochondrial m^1^A modification.

In this respect, ModiDeC should not be considered a competitor to approaches that perform global scans for a specific modification but as a highly precise, complementary tool for accurately determining and validating certain combinations of motif and modification, with a biological and/or clinical context. Its strength is its ability to determine the exact position and type of a modification, especially if there is great uncertainty about whether a modification even exists in a particular region. Using validation data with known ground-truth and unmodified IVT data, we showed that ModiDeC could correctly identify the type of modification with 100% accuracy, even if different modifications at different locations were to be detected in the same sample or even within the same motif (see Fig. 1e; we added all four modifications once each at the same position in the DRACH motif “AGACA”).

ModiDeC clearly distinguishes itself from other excellent tools already available for detecting RNA modifications that have been trained, for example, with Randomer Oligonucleotides “Randomers” (as e.g. in the case of the Dorado base callers) and are optimized for large-scale screenings for only one particular type of modification. Although methods trained with randomers offer great advantage if the aim is to find as many modifications as possible to obtain an overview of the entire epitranscriptomic landscape, they are less favorable if one is interested in a specific motif and wants to achieve the highest possible accuracy in terms of position and quantification. Randomly generated sequences have the limitation (purely statistically) that some motifs will occur particularly frequently in the training data and will, therefore, be recognized particularly well, while other motifs will be encountered relatively rarely in the training data and will, therefore, be learned particularly poorly. However, by using randomer-based approaches, it is very challenging and demanding to influence which motifs are learned particularly well. It cannot be assumed that exceptionally well-trained motifs are biologically significant; indeed, underrepresented motifs might be precisely those that are of particular interest to the application-oriented user.

As also apparent with our own training data, even under identical conditions, and with a very large amount of training data (30,000 identical sequences per motif– modification combination), some motifs were harder to learn and already showed a high false-negative rate on the synthetic data, whereas other motifs were much easier to remember (see Fig. 1e and Supplementary Fig. 4). If such a “difficult” motif is then also underrepresented in randomized training data, it will most likely have a poor detection rate when the model is applied to biological data.

In a further complication, false-positive predictions, false negatives, and misclassified RNA modifications for transcriptome-wide predictions can rarely be validated owing to a lack of gold-standard data sets. For m^6^A, larger experimentally validated benchmarking data sets for HEK293T cell lines already exist^30,50,64^, which allows predictions about modifications to be validated to a certain extent. Also, a reference catalog has been recently published for pseudouridine^41,59^. However, these experimentally validated databases should not be regarded as gold standards either, as false-positive or false-negative predictions cannot be ruled out, as exemplified by several contradictory statements regarding the existence of certain modifications. Even for the already very well-familiar m^6^A landscape of HEK293T cell lines, considerable uncertainty exists, as shown by the overlap between the CHEUI base-caller data and the GLORI database, which was tested in two independent studies in 2024^47,50^. Chan et al. reported a site-level overlap of 0.64 and Mateos et al. determined it to be 0.85. Similar observations have been made for pseudouridine sites^66^ and were also currently reported with respect to m^1^A sites^19,65^. Furthermore, a considerably tissue-specific and even time-dependent modification landscape must be assumed, which makes validation in other tissues or cell types highly speculative.

In this context, a tool such as ModiDeC can be of great utility, as it can be applied explicitly to ill-defined and/or difficult-to-detect modifications and motifs for validation. In cases for which we knew the exact truth and were thus able to validate the predictions (Figs. 1–5), ModiDeC correctly recognized the presence of a modification in one nucleotide every time, correctly identified the type of modification each time, and also recognized the exact position of the modification in the validation data, even if the sequence surrounding the motif was completely new because the algorithm had been trained on different training data with a different sequence context. As demonstrated in several of the biological samples, we were able to show that when interested in specific issues or regions, it is not necessary to first generate all possible motifs and permutations around a particular modification. Rather, we have already achieved very significant performance with a very limited number of motifs (10 per modification).

Peak detection was made further reliable by continually training and validating our findings with matching IVT data that had a negligible false-positive rate for the annotated positions (< 2%) and a low false-positive rate on the overall IVT transcripts (< 5% false positives in >97% of the cases). A result of particular note is that ModiDeC seems to have versatility in identifying motifs that differ from those used for training but contain variations in the sequence (see Supplementary Table 4). This indicates that ModiDeC has potential for identifying features that have not been explicitly learned from the training pool and would thus also be suitable for use in human data, in which a large number of genetic variants, such as single nucleotide polymorphisms, must be assumed. We also see ModiDeC as particularly suitable for predicting unknown modification sites, which can then be experimentally validated. As shown in Fig. 5 and the Supplementary Information (SI Figure 8), instances arose where ModiDeC predicted Gm or inosine that could not be validated due to a lack of reference data in the literature, but which due to the experimental setup (high predictions frequency for the blood samples, with simultaneous correctly recognized negative IVT) are interesting candidates for further studies.

A major unresolved issue arises from a systematically comparing models that predict modifications from DRS data for use in addressing biological questions. Most of the models already developed have been trained on RNA002 chemistry and only a few on data from RNA004 sequencing. Therefore, only a few benchmarks are representative, as these require that models are compared on differently generated data (some on RNA002 and some on RNA04). Additionally, all would have the problem that an exact validation, apart from synthetic oligos, would be impossible due to the lack of a gold standard for biological data. Since such benchmarking would have been outside the scope of this study—especially for the many modifications considered here—we have refrained from doing so; instead we refer to a recently published systematic review for m^6^A base caller^67^.

Furthermore, we call for an extensive and coordinated community effort to provide ground-truth data for establishing a gold-standard database for various tissue types, to better validate *in silico* predictions and, thus, to better train and compare the various tools for predicting modification detection.

Together with ModiDeC, we offer all our data as training and validation data for other tools and classifiers. We are providing an extensive database of carefully designed high-quality training and validation data for various RNA modifications, together with matching IVT data for synthetic data, for HEK293 cell lines and also for human peripheral blood samples. Each sample was sequenced once with SQK-RNA002 and once with SQK-RNA004 so that even community models that were designed for older chemistry can be retrained for special purposes with our data and to optimize them for specific motifs.

Although ModiDeC can be used in various scenarios including human peripheral blood, our neural network model is limited in that only 40 motifs were generated for the initial model training. This limits the tool’s utility in comprehensive genome-wide screens for routine diagnostic purposes despite its ability to extract features outside the training pool. To offer the possibility of personalized training for individual research questions, we developed a GUI that allows the user to retrain ModiDeC to search for user-defined specific sequences or motifs. We also see great potential in that our data pool will be a seed for further community-driven data generation. With the help of the GUI, any lab interested in a specific region in the transcriptome of a species can easily order matching oligos and train ModiDec on this position or the corresponding motif, and then make it available to the community.

We argue that the ease of extension and customization of ModiDeCs by anyone makes it a valuable tool for studies on RNA modification in biology and medicine. Also, if combined with models optimized for genome-wide screenings, ModiDeC could be a complementary high-precision validation tool.

## Methods

### Plasmid preparation and *in vitro*-transcription (IVT)

Plasmid sequences were cloned in a pUC57 vector containing an internal T7 promotor, followed by the desired template sequence and a restriction enzyme cutting site (either MscI or SacII) at the end. The linearization reaction was performed overnight following the respective restriction enzyme manufacturer’s instructions (ThermoFisher Scientific). Linear plasmids were purified by phenol–chloroform extraction followed by ethanol precipitation. Linearization of the plasmids and their overall quality was confirmed by separation on an agarose gel and quantified using a spectrophotomer (NanoDrop One).

IVT was performed using a HiScribe T7 High Yield RNA Synthesis Kit according to the manufacturer’s instructions. If the transcription product required 5’ phosphorylation for ligation, the concentration of the terminal NTP (GTP, ATP, CTP, or UTP) was reduced and the reaction supplemented with NMP (GMP, AMP, CMP, or UMP) at a ratio of 1:5 NTP/NMP.

Transcription products were purified using a Monarch RNA Cleanup Kit (New England Biolabs, T2040), and their quality was assessed by separation on agarose gel and NanoDrop One analysis.

### Splinted ligation of RNA constructs

RNA oligos were 5′-phosphorylated to utilize them in the ligation reaction. Phosphorylation was performed using T4 polynucleotide kinase (New England Biolabs, M0201) following manufacturer’s instructions. Phosphorylated oligos were purified using Oligo Clean & Concentrator Kit (Zymo research, D4060) according to the manufacturer’s instructions.

Ligations were performed using T4 RNA ligase 2 (New England Biolabs, M0239), combining equimolar amounts of *in vitro*-transcribed RNA and the ligating oligo, 1× T4 RNA ligase buffer, 10% w/v PEG-8000, 10 U T4 RNA ligase 2, as well as a complementary cDNA splint ensuring the correct order of ligation. Here, 2% less cDNA splint was supplied compared to the amount of *in vitro*-transcribed RNA or oligos present.

The reaction mixture was prepared as above, omitting T4 RNA ligase 2 for an initial denaturation step at 75°C. After cooling to 25°C over 15 min, T4 RNA ligase 2 was introduced and the reaction was incubated at 16°C overnight.

The ligation reaction was stopped by digestion of the cDNA splint using DNase I (ThermoFisher Scientific, EN0525) according to the manufacturer’s instructions, followed by purification using an Oligo Clean & Concentrator Kit.

The purified ligated constructs were polyadenylated using *E. coli* poly(A) polymerase (New England Biolabs, M0276) according to the manufacturer’s instructions, and purified again using an Oligo Clean & Concentrator Kit before proceeding to library preparation. Purity and concentration after each purification step were assessed using a NanoDrop One spectrophotometer.

### Isolation of nuclear and cytoplasmic rRNA

Isolation of nuclear and cytoplasmic rRNA was performed as follows: Cells were trypsinized, washed with cold phosphate-buffered saline, and resuspended in nuclei isolation buffer (NIB; 10 mM Tris-HCl pH 7.4, 10 mM NaCl, 3 mM MgCl₂, 0.1% IGEPAL CA-630, 0.1% Tween-20, 1% BSA, 0.15 mM spermine, 0.15 mM spermidine, and 0.2 U/μl RNase inhibitor). Homogenization was performed with a loose pestle (10 strokes), followed by incubation for 15 min on ice and then centrifugation at 1190 x *g* for 5 min at 4°C. The cytoplasmic fraction (supernatant) was collected, and Trizol was added at RT. The remaining pellet was washed twice with 500 μl of ice-cold NIB, centrifuged, and resuspended in 300 μl of ice-cold NIB. For isolation of nuclei, 300 μl of 50% Optiprep solution (Stemcell Technologies, 07820) was mixed with the homogenate to form a 25% Optiprep mix. This was layered over 30% and 40% Optiprep gradients in a 2 ml tube, centrifuged at 20800 x *g* for 20 min at 4°C, and nuclei (∼600 μl) were collected from the 30%–40% interface. The nuclei were washed with nuclei wash buffer (NWB; 10 mM Tris-HCl pH 7.4, 10 mM NaCl, 3 mM MgCl₂, 0.1% Tween-20, 1% BSA, and 0.2 U/μl RNase inhibitor), centrifuged at 1635 x *g* for 10 min at 4 °C, and the supernatant was removed. Nuclease-free water was added the nuclei to a volume of 500 μl and 500 μl of Trizol was added prior to RNA isolation.

For RNA isolation, nuclear and cytoplasmic fractions (∼1 mL) were incubated at RT for at least 5 min. Chloroform (200 µl) was added to the nuclear and cytoplasmic fractions. Samples were vortexed for 15 s, incubated at RT for 3 min and centrifuged for 15 min at 20800 x *g* at 4°C. The upper aqueous phase (∼550 µl) was transferred into a fresh Eppi and 500 µl of isopropanol was added, thoroughly resuspended and incubated at RT for 15 min. Sample were centrifuged for 10 min at 20800 x *g* at 4°C. The supernatant was discarded and the pellet was washed with 75% ethanol in nuclease-free water, followed by centrifugation for 5 min at 7500 × *g* at 4°C. The supernatant was discarded, and pellet was air dried for 5–10 min. The RNA pellet was dissolved in 30 µl of RNAse-free water and mixed in a high-shear rotor-stator sample mixer (Hula Mixer, ThermoFisher Scientific) for 10 min at RT and RNA concentration was quantified using the Qubit dsDNA Assay Kit (Invitrogen, Q32851).

### Direct RNA sequencing library preparation

Libraries were prepared with 1 µg of polyadenylated input material using either SQK-RNA002 or SQK-RNA004 kits (Oxford Nanopore Technologies) according to the manufacturer’s standardized protocol. The prepared libraries were quantified using the Qubit dsDNA Assay Kit (Invitrogen, Q32851). Libraries were loaded onto MinION flow cells and sequenced with MinKNOW (Version 24.02.26) set to fast live base calling until no more sequencing pores were available.

### Peripheral blood

Peripheral blood was obtained from a healthy volunteer. RNA was extracted using a PAXgene Blood miRNA Kit (Qiagen) according to the manufacturer’s protocol, except that the RNA was eluted in nuclease-free water instead of the provided buffer. The RNA was characterized using the Bioanalyzer (Agilent) total RNA nano assay according to the manufacturer’s protocol. The RNA had a concentration of 363 ng/μl and an RNA integrity number (RIN) of 7.2. Globin mRNA was depleted with the GLOBINclear-Human Kit (Thermofisher Scientific, AM1980) according to the manufacturer’s protocol; this was performed four times on a total input amount of 20 μg of RNA to yield 11 μg of globin-mRNA-depleted RNA. The concentration was measured using a Qubit RNA HS Assay (Thermofisher Scientific). Two micrograms of RNA was stored for later use in the direct RNA run. The remaining 9µg of RNA were split into three aliquots of 3µg respectively used for poly(A) selection with the NEBNext Poly(A) mRNA Magnetic Isolation Module (New England Biolabs) according to the manufacturer’s protocol; The poly(A) selection resulted in a total yield of 23 ng, as measured with the aforementioned Bioanalyzer assay. The sample had an average length of ∼1 kb. The sample concentration was measured again using the aforementioned Qubit assay. Reverse transcription, PCR, *in vitro* transcription (IVT), polyadenylation and 5’ capping was carried out according to the methods of Tavakoli et al. (2023) with modifications as follows. Each IVT primer was used in the PCR at a final concentration of 0.5 μM. The input amount of mRNA used for reverse transcription and PCR was 7.1 ng. The output amount of cDNA was 905 ng, as measured with a Qubit DNA HS Assay (Thermofisher Scientific). IVT was performed twice, each with an input of 126.7 ng of cDNA. The outputs were pooled to yield 4.9 μg of RNA, as measured with the Qubit RNA HS assay. Libraries were prepared using the direct RNA sequencing Kit (SQK-RNA004, Oxford Nanopore Technologies). The library output was 167 ng of RNA/cDNA hybrid (Qubit DNA HS Assay). The complete library was loaded onto the PromethION RNA Flow Cell (FLO-PRO004RA).

### Neural network training and data processing

ModiDeC is a two-input neural network that combines personalized and new structured inception blocks with LSTM to classify RNA modifications. To train the neural network, raw signals and the respective one-hot encoded sequences are necessary. A minimal number of passages are necessary for the neural network to obtain the correct training outcome. The raw pod5 files from all sequencing runs were first base-called using the high-accuracy model in Dorado v0.7.3. Next, the raw signal was pre-processed by squiggling the raw signal using Remora (Oxford Nanopore Technologies), assigning the corresponding reference in one-hot encoding, and splitting both raw signals into chunks (training input). We used a chunk size of 400 time points (input 1) for the raw signal and max sequence length of 40 nucleotides (input 2). In parallel, the one-hot encoding output of the neural network is also generated, in which each base is converted into its respective modified variant or into an unmodified label. A mixture of modified and unmodified data was essential for training ModiDeC, including IVT data for training the neural network. Ultimately, the neural network was trained for three epochs with a learning rate of 0.0001 using the Adam optimizer.

To generate the training data for ModiDeC, the “ModiDeC data curation” GUI was used to pre-process 30,000 reads for each motif used in this work (Fig. 1e). Ten thousand high-quality IVT reads (selected using samtools and filter flag q=20) that aligned to the gen-code v46 transcripts were also pre-processed using the GUI and then used for the training. For the sample analysis we established a minimum threshold of 500 reads for each transcript to ensure statistical reliability. This value was decreased to 80 for IVT data as the number of reads produced during IVT measurement can be small.

## Code Availability

The neural network and the GUI codes for ModiDeC are available at GitHub: https://github.com/mem3nto0/ModiDeC-RNA-modification-classifier, where tutorials for retraining and/or using ModiDeC are also provided.

## Data Availability

All data for this study will be made freely available to the community once the manuscript has been conditionally accepted.

## Acknowledgements

This work was partly funded by the Deutsche Forschungsgemeinschaft (DFG, German Research Foundation; project no. 439669440 TRR319 RMaP (TP A05/C01/C03 to M.H.and S.M) and seed funding from TP Z (S.G. and K.F). N.A. and S.G. acknowledge funding from the Forschungsinitiative Rheinland-Pfalz and the ReALity initiative of the Johannes Gutenberg University Mainz. S.G. and L.L. acknowledge funding from the Boehringer Ingelheim Stiftung. “K.F. acknowledges Mitoscience, Ministerium für Wirtschaft, Verkehr, Landwirtschaft und Weinbau Rlp“

