## Supplemental Information for "ModiDeC: a multi-RNA modification classifier for direct nanopore sequencing"

**Supplementary Information**

### SI 1. Creation of training data sets for ModiDeC

**Supplementary Table 1:** List of all oligonucleotides used for training ModiDeC. The incorporated modified bases **pseudouridine (Ψ), 2'-*O*-methylguanosine (Gm), *N*^6^-methyladenosine (m^6^A), inosine (I)** and ***N*^1^-methyladenosine (m^1^A)** are highlighted and color-coded as they appear in the main text and supplementary section.

| **Name** | **Full sequence** | **Internal modification** |
| --- | --- | --- |
| unmod. | GAUACGGGAGACAGCCACCGGAAUACGGGAGACAGCCACCUC | none |
| Psi #1 | GAUACGGGAGUCAGCCACCGGAAUACGGGAGΨCAGCCACCUC | Pseudouridine (Ψ) |
| Psi #2 | GAGCCAUGAUUAAGAGGGAGGAAGCCAUGAUΨAAGAGGGAUC | Pseudouridine (Ψ) |
| Psi #3 | GAGGUGACUCUAGAUAACCGGAAGGUGACUCΨAGAUAACCUC | Pseudouridine (Ψ) |
| Psi #4 | GAACCACGGGUGACGGGGAGGAAACCACGGGΨGACGGGGAUC | Pseudouridine (Ψ) |
| Psi #5 | GAUGCGUGCAUUUAUCAGAGGAAUGCGUGCAΨUUAUCAGAUC | Pseudouridine (Ψ) |
| Psi #6 | GAAGGCCAAGUAGCCCGUGGGAAAGGCCAAGΨAGCCCGUGUC | Pseudouridine (Ψ) |
| Psi #7 | GAAUGCCACCUAGGGCAAGGGAAAUGCCACCΨAGGGCAAGUC | Pseudouridine (Ψ) |
| Psi #8 | GAAAUGAUUCUUAGCAGGGGGAAAAUGAUUCΨUAGCAGGGUC | Pseudouridine (Ψ) |
| Psi #9 | GACCUAGGGAUCUUCCGAAGGAACCUAGGGAΨCUUCCGAAUC | Pseudouridine (Ψ) |
| Psi #10 | GAUCGAGGUCUCUGCAACCGGAAUCGAGGUCΨCUGCAACCUC | Pseudouridine (Ψ) |
| Gm #1 | GAUACGGGAGGCAGCCACCGGAAUACGGGAGGmCAGCCACCUC | 2'-*O*-Methylguanosine (Gm) |
| Gm #2 | GACCGGGGAGGUAGUGACGGGAACCGGGGAGGmUAGUGACGUC | 2'-*O*-Methylguanosine (Gm) |
| Gm #3 | GAUAAUAACGGGUCUGUGAGGAAUAAUAACGGmGUCUGUGAUC | 2'-*O*-Methylguanosine (Gm) |
| Gm #4 | GACGUAGUUGGAUCUUGGGGGAACGUAGUUGGmAUCUUGGGUC | 2'-*O*-Methylguanosine (Gm) |
| Gm #5 | GAACUUCUUAGAGGGACAAGGAAACUUCUUAGmAGGGACAAUC | 2'-*O*-Methylguanosine (Gm) |
| Gm #6 | GAUGUGAAUUGCAGGACACGGAAUGUGAAUUGmCAGGACACUC | 2'-*O*-Methylguanosine (Gm) |
| Gm #7 | GACAUUGGAGGGCAAGUCUGGAACAUUGGAGGmGCAAGUCUUC | 2'-*O*-Methylguanosine (Gm) |
| Gm #8 | GACUUAGCAUGCGAGAGGUGGAACUUAGCAUGmCGAGAGGUUC | 2'-*O*-Methylguanosine (Gm) |
| Gm #9 | GAAAUCUUUCGCCUUUUACGGAAAAUCUUUCGmCCUUUUACUC | 2'-*O*-Methylguanosine (Gm) |
| Gm #10 | GAGGACUAGAGGCCGUAUGGGAAGGACUAGAGmGCCGUAUGUC | 2'-*O*-Methylguanosine (Gm) |
| m^6^A #1 | GAUACGGGAGACAGCCACCGGAAUACGGGAGm^6^ACAGCCACCUC | *N*^6^-Methyladenosine (m^6^A) |
| m^6^A #2 | GAGUGCCAGGACCGACCAUGGAAGUGCCAGGm^6^ACCGACCAUUC | *N*^6^-Methyladenosine (m^6^A) |
| m^6^A #3 | GACACCAGUGACUCCCAUAGGAACACCAGUGm^6^ACUCCCAUAUC | *N*^6^-Methyladenosine (m^6^A) |
| m^6^A #4 | GAUAAACGAGACCGUCUAGGGAAUAAACGAGm^6^ACCGUCUAGUC | *N*^6^-Methyladenosine (m^6^A) |
| m^6^A #5 | GAGUUAAGAGACUGAAUCUGGAAGUUAAGAGm^6^ACUGAAUCUUC | *N*^6^-Methyladenosine (m^6^A) |
| m^6^A #6 | GAAGUACAAAACAAUCAUUGGAAAGUACAAAm^6^ACAAUCAUUUC | *N*^6^-Methyladenosine (m^6^A) |
| m^6^A #7 | GAUUCAGAGAACCACUUGAGGAAUUCAGAGAm^6^ACCACUUGAUC | *N*^6^-Methyladenosine (m^6^A) |
| m^6^A #8 | GAUAAACUUAACUCCAAAAGGAAUAAACUUAm^6^ACUCCAAAAUC | *N*^6^-Methyladenosine (m^6^A) |
| m^6^A #9 | GAUCACUCGAACUUCAAGCGGAAUCACUCGAm^6^ACUUCAAGCUC | *N*^6^-Methyladenosine (m^6^A) |
| m^6^A #10 | GAUUGUGGUAACGUCCCCAGGAAUUGUGGUAm^6^ACGUCCCCAUC | *N*^6^-Methyladenosine (m^6^A) |
| Inosine #1 | GAUACGGGAGACAGCCACCGGAAUACGGGAGICAGCCACCUC | Inosine (I) |
| Inosine #2 | GAGCUCGCUUAGCAUGCGAGGAAGCUCGCUUIGCAUGCGAUC | Inosine (I) |
| Inosine #3 | GAGAGGGUCGAGAUCCACUGGAAGAGGGUCGIGAUCCACUUC | Inosine (I) |
| Inosine #4 | GAUUGCGGAUACCGAACGAGGAAUUGCGGAUICCGAACGAUC | Inosine (I) |
| Inosine #5 | GAGAGCUGGCACGAGGCUCGGAAGAGCUGGCICGAGGCUCUC | Inosine (I) |
| Inosine #6 | GAGCCCCAAAACGGCCGAUGGAAGCCCCAAAICGGCCGAUUC | Inosine (I) |
| Inosine #7 | GAUGGAGAGCACGAGGCCCGGAAUGGAGAGCICGAGGCCCUC | Inosine (I) |
| Inosine #8 | GAAUCUUGUCAAUGUCGUCGGAAAUCUUGUCIAUGUCGUCUC | Inosine (I) |
| Inosine #9 | GAUAUACGUCAGGCACCCUGGAAUAUACGUCIGGCACCCUUC | Inosine (I) |
| Inosine #10 | GAGGAACGGCACAGUUAUUGGAAGGAACGGCICAGUUAUUUC | Inosine (I) |
| m^1^A #1 | CCACm^1^AAACCAUGGUGAGCAA | *N*^1^-Methyladenosine (m^1^A) |
| m^1^A #2 | AACGCCTGGCm^1^AGCCGGAAGCC | *N*^1^-Methyladenosine (m^1^A) |
| m^1^A #3 | AACGCCTGGCAGCCGGAAGCC | none |

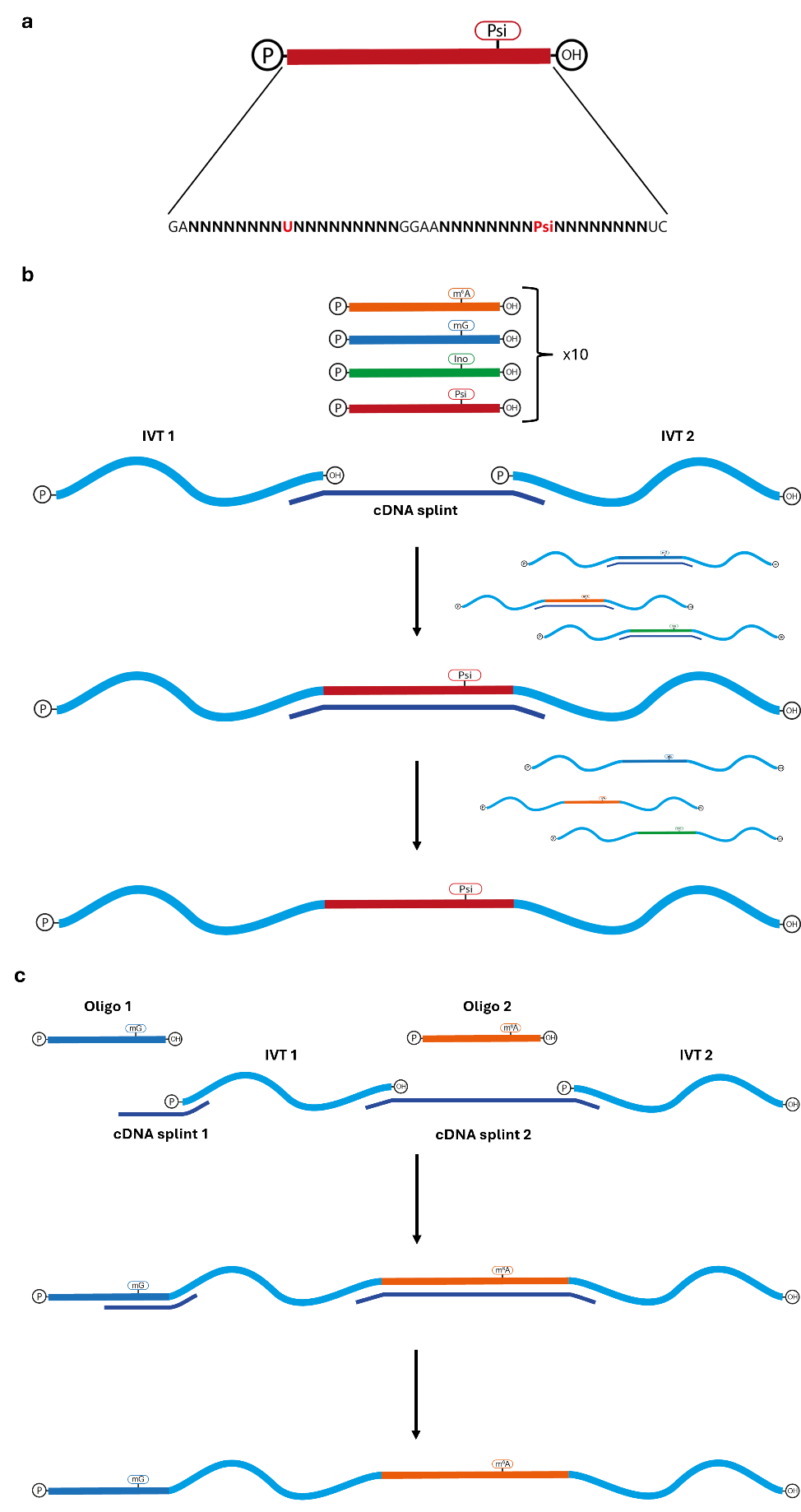

**Supplementary Fig. 1:** **Oligonucleotide design and incorporation of oligonucleotides into sequencing construct via splinted ligation**. (a) Oligonucleotides contain the modified and unmodified nucleotide (red) flanked by a 9-mer sequence of choice (bold, black).
(b) Oligonucleotides are incorporated into a larger RNA construct via splinted ligation. A cDNA splint complementary to the 5' and 3'- ends of the IVT RNA and the oligonucleotides aligns the components for ligation by T4 RNA Ligase 2. Following ligation, the cDNA splint is digested by DNase, leaving behind the ligated product. (c) Ligation of sequencing constructs containing multiple oligonucleotides. Two complementary cDNA splints are supplied, one for each oligonucleotide. The alignment and DNase digestion process is as described in panel (b).

### SI 2. Testing neural network efficiency over reads used for the training phase

In this part of the study we investigated whether we could identify a certain number of reads to use for the training phase. To this end, we trained ModiDeC to analyze pseudouridine RNA modifications (Ψ) using different amounts of training data. Specifically, we used 1,000, 5,000, 10,000, 20,000, and 30,000 reads for the training phase and then tested the efficiency of the neural network on new constructs (different from the training constructs) that contain a Ψ modification at a specific position. An unmodified motif is also present in the testing construct and used for the analysis. The confusion matrices for 1,000, 5,000, 10,000, 20,000, and 30,000 reads (Supplementary Fig. 2a–e) show prediction versus expectation for the modified position (Ψ) and for the unmodified position (G). We observed that increasing the number of reads for the training increased the percentage of correctly called Ψ (low right squares on the heatmaps), reaching a maximum value for 30,000 reads (Supplementary Fig. 2e). The unmodified position seems to follow a different trend than the modified one. To better understand the global effect of data training of neural network performance, we averaged the accuracy values of the main diagonal for each heatmap and then plotted the results versus the number of reads (Supplementary Fig. 2f). This plot shows that general performance depends on the number of training data used. We find the curve somewhat saturated at 10,000 reads, which suggests using a certain minimum number of reads in the training phase. However, it is difficult to reach a general conclusion because the results presented in Fig. 1e and Supplementary Fig. 4 show different frequencies of modification detection that depend on the RNA motif, although uniform training data were used during the training phase (each motif was trained with approximately 30,000 reads). Despite these mixed results, we suggest using minimum 10,000 reads as a reference value for future training.

**Supplementary Fig. 2:** Confusion matrices on test data using ModiDeC trained with 1,000 (a), 5,000 (b), 10,000 (c), 20,000 (d) and 30,000 (e) reads. Decimal fraction values are written inside each square. It is possible to observe an increase of true positives for pseudouridine predictions by increasing the training data batch. f) True-positive average values (y axis) plotted against the size of the training data batch (x axis). True positive average values were calculated by averaging the main diagonal of each confusion matrix. Performance seems to saturate after 10,000 training data.

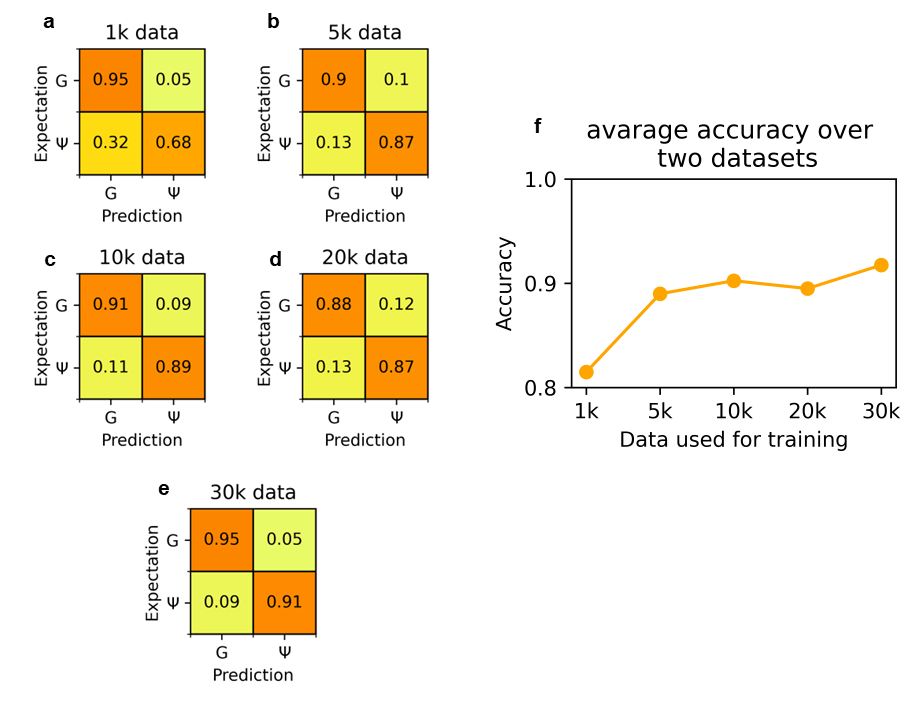

**Supplementary Table 2:** Modified 9-mer sequences for the 40 oligos analyzed in Fig. 1e measured with the 004 kit. The table shows the true-positive, false-negative and modification miscall percentages obtained using ModiDeC. The expected true-positive percentage is 100% for all the 9-mers.

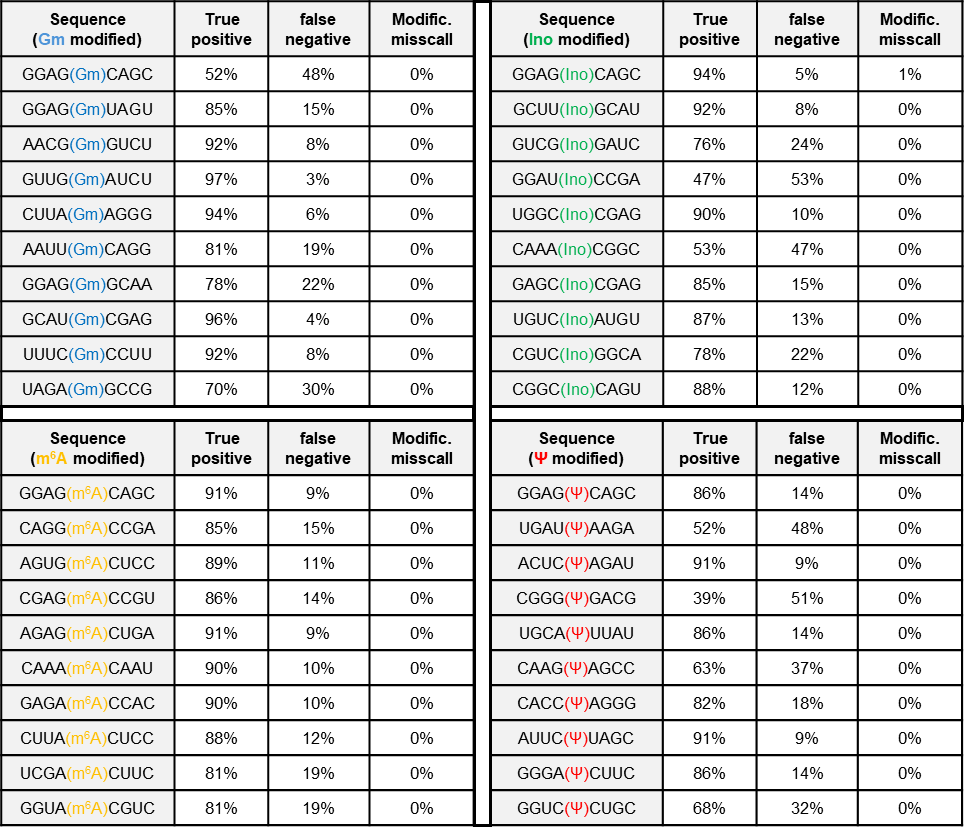

### SI 3. ModiDeC validation results on unmodified read region in 004 ONT kit

ModiDeC results for the SQK-RNA004 data for unmodified reads region. The sequence region is the same for modified (Fig. 1e) and unmodified (Supplementary Fig. 3) data.

**Supplementary Fig. 3:** The modification frequency on the test data for the 40 motifs used in this work for unmodified reads for (SQK-RNA004 kit). The green letter shows the canonical letter that is modified in Fig. 1e. The unmodified region shows baseline results as expected.

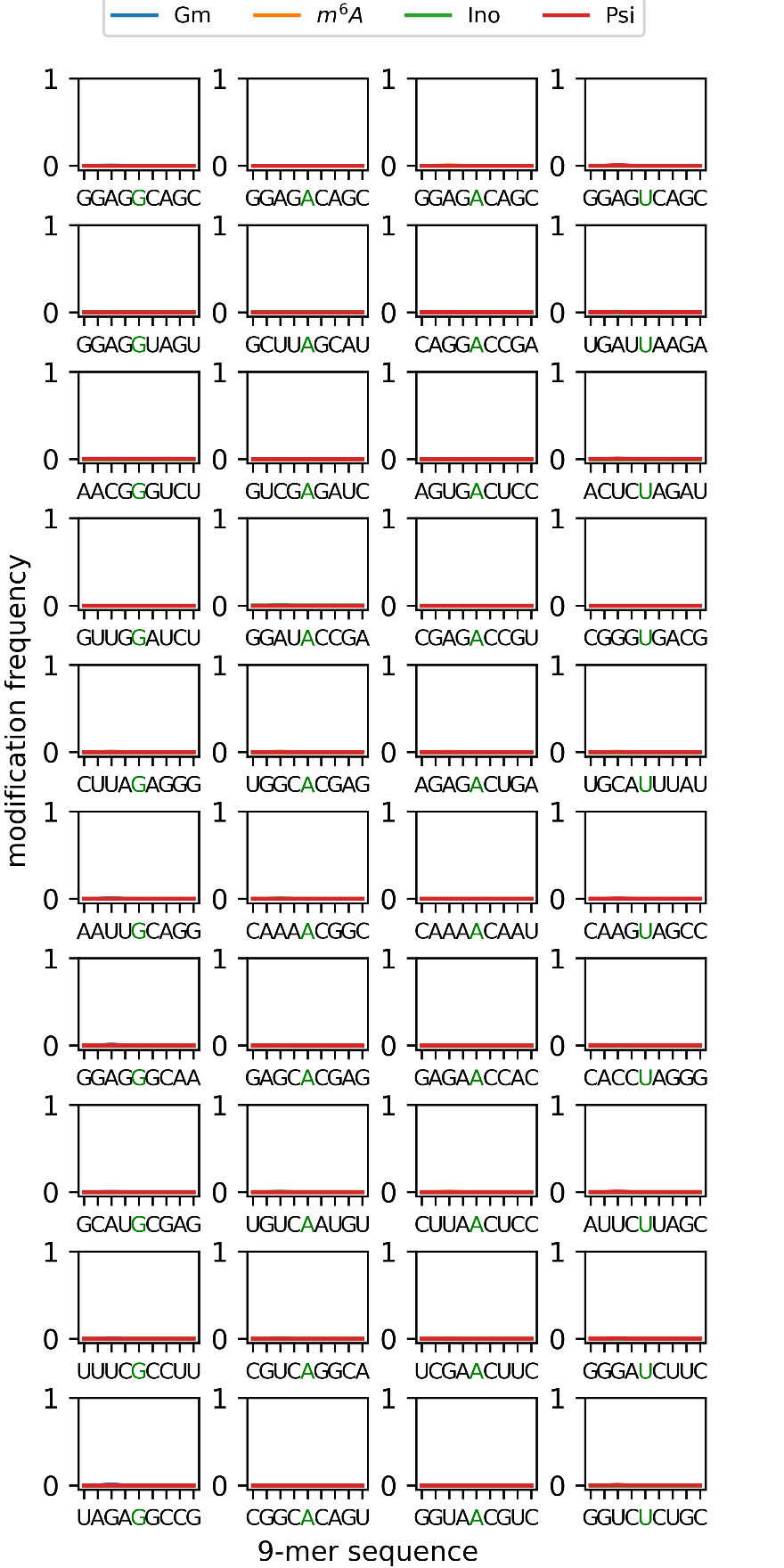

### SI 4. ModiDeC validation results for SQK-RNA002

Here we show the results obtained from the validation of ModiDeC after training the neural network on the SQK-RNA002 Kit. A comparison of the data in Fig. 1e and Supplementary Fig. 4 shows the frequency of detecting modifications is similar for the 002 and 004 kits. Supplementary Fig. 5 shows the results for the unmodified region. As expected, this region gave a practically baseline result, indicating that reads were not modified in the designed area.

**Supplementary Fig. 4:** The modification frequency on the test data for the 40 motifs used in this work (SQK-RNA002 kit). The red letter shows the modified base. We obtained similar results with the SQK-RNA004 kit data (see Fig. 1e).

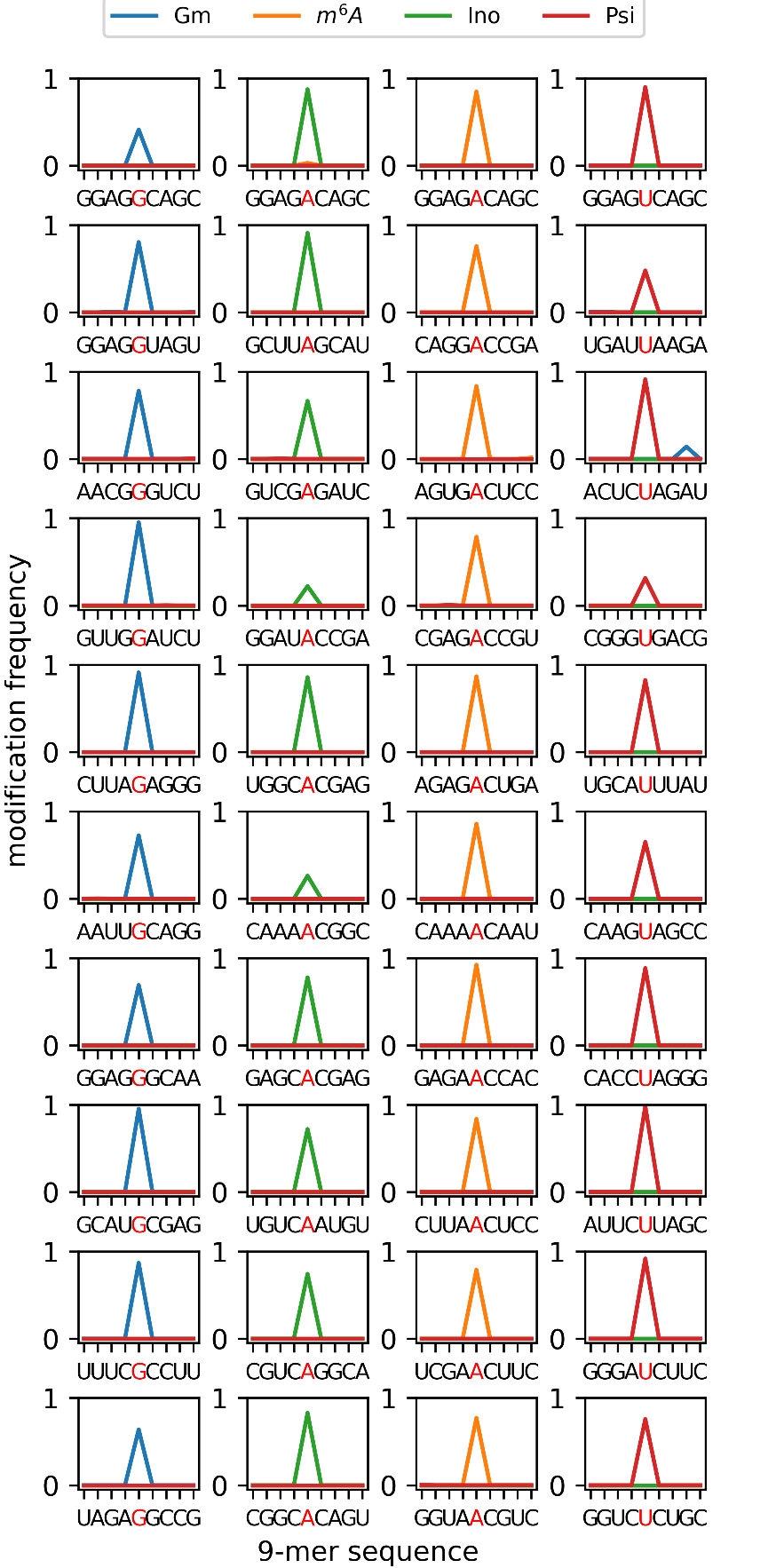

**Supplementary Fig. 5:** The modification frequency on the test data for the 40 motifs used in this work for unmodified reads with the SQK-RNA002 kit. The green letter shows the canonical letter that is modified in Fig. 1e. The unmodified region shows baseline results as expected.

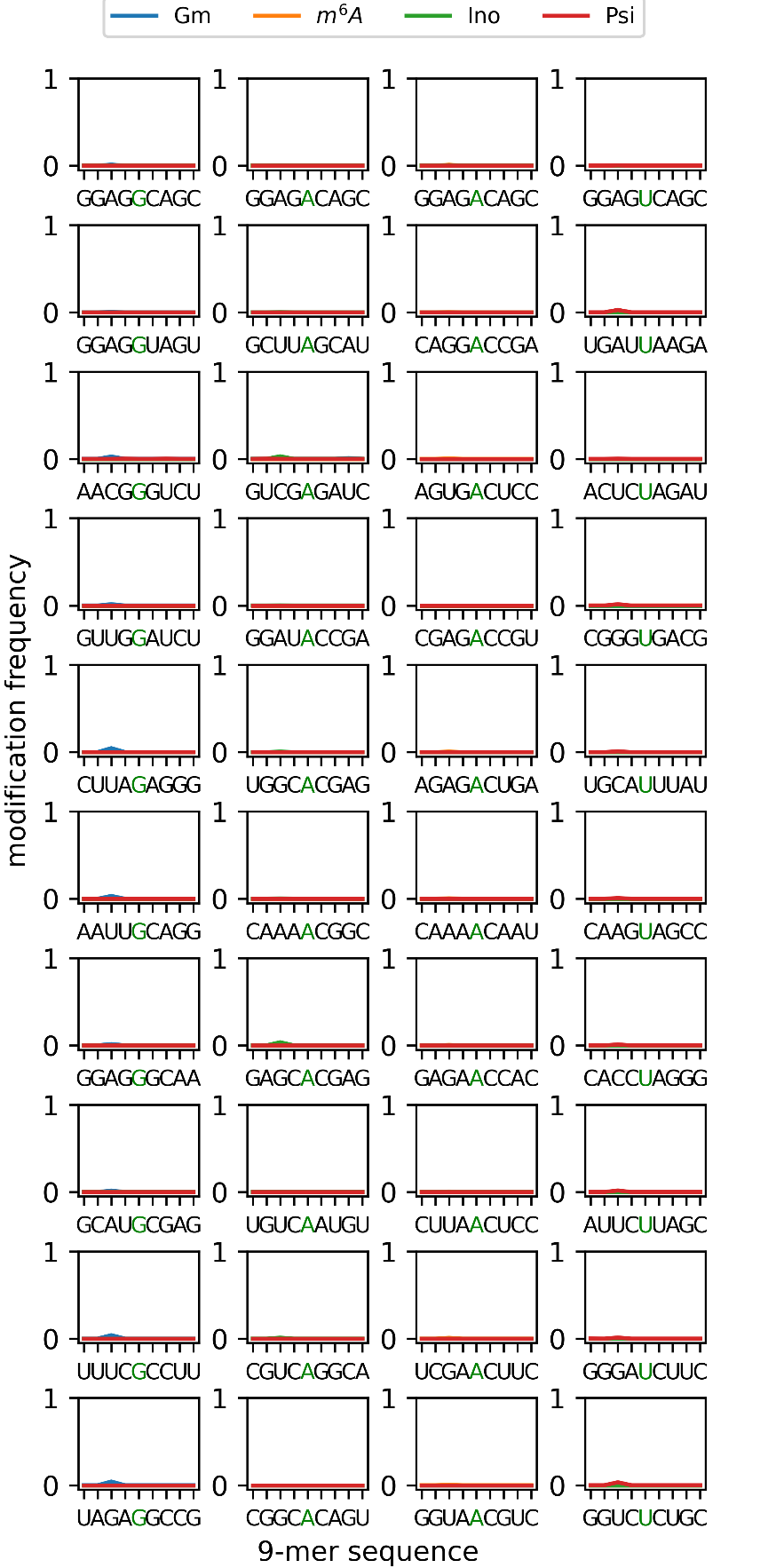

.

**Supplementary Table 3:** Modified 9-mer sequence for the 40 oligos analyzed in Fig. 1e measured with the SQK-RNA002 kit. The table shows the true positive, false negative and modification miscalls obtained using ModiDeC for the oligos analysis. Results are shown for each modified 9-mer. The expected true positive value is 100% for all the 9-mers.

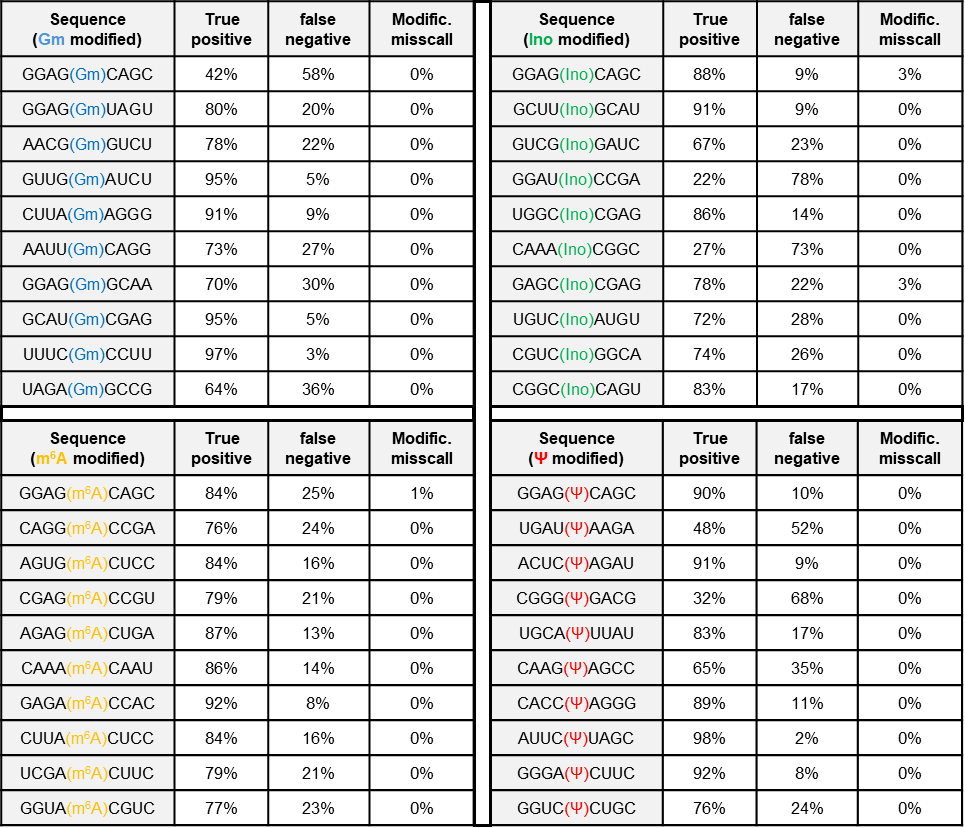

### SI 5. Analysis of HEK293T cells with Dorado: selective organelle and ribosomal RNA

We performed Dorado analysis on HEK293T cells containing orthogonally translating film-like organelles. In details, EGFP and mCherry fluorescence protein sequences were analyzed using Dorado to quantify the amount of pseudouridine in specific sequence positions. Supplementary Fig. 6 shows the modification frequencies of pseudouridine (Psi), U–C mismatch, and unmodified reads detected by the Dorado model (Oxford Nanopore Technology) for the following three systems: without organelle (nO), with organelle (O), and control (C). The expected reference position of modification is shown below each bar plot (114 for EGFP and 564 for mCherry). Tables containing the results of the Dorado analysis for the three samples are shown next to each bar plot in Supplementary Fig. 6.

**Supplementary Fig. 6:** Dorado analysis on EGFP (a) and mCherry (b) sequences for systems without organelle (nO), with organelle (O), and control (C). On the right of the plot, the Dorado analysis values for the three samples are tabulated.

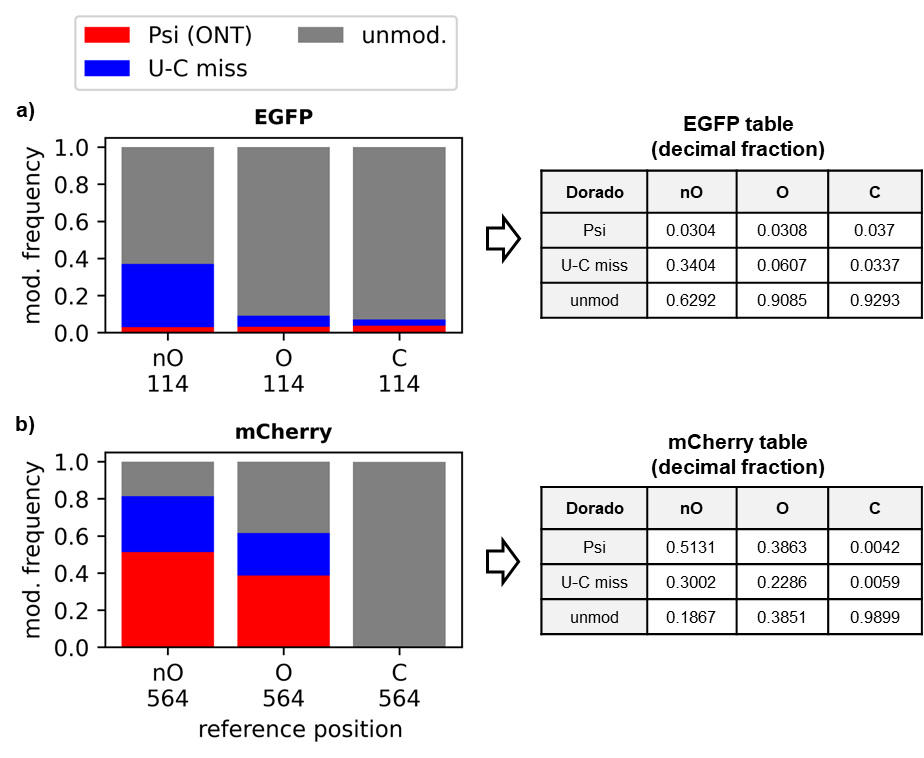

Additional information about the ModiDeC analysis on ribosomal RNA are reported below in the Supplementary Table 4.

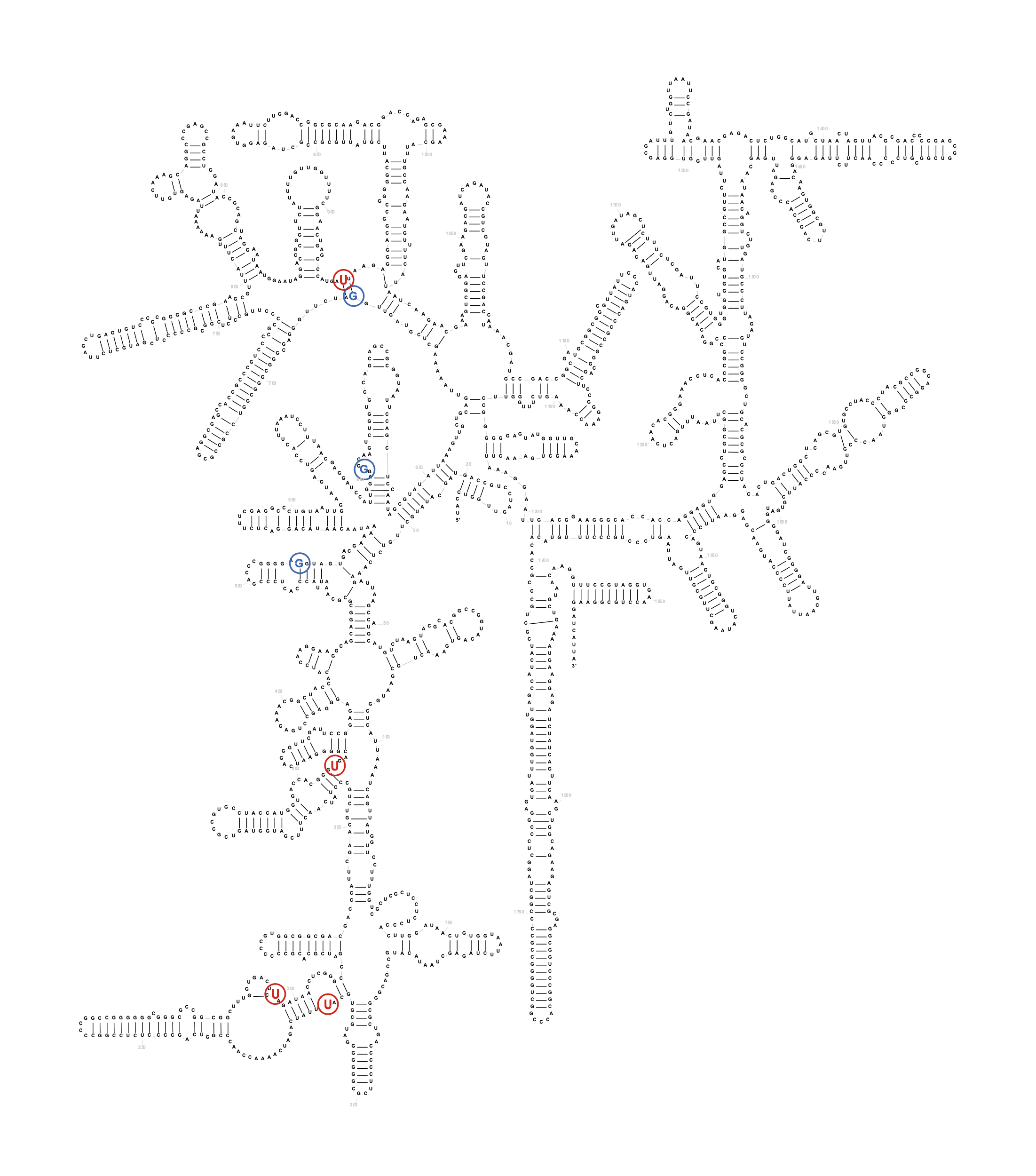

**Supplementary Fig. 7:** RNA45SN1 RNA complete structure. The letters highlighted are the modified bases with trainings-motives as recognized by ModiDeC (red = Psi, blue = Gm). The numbers next to these letters indicate the reference position.

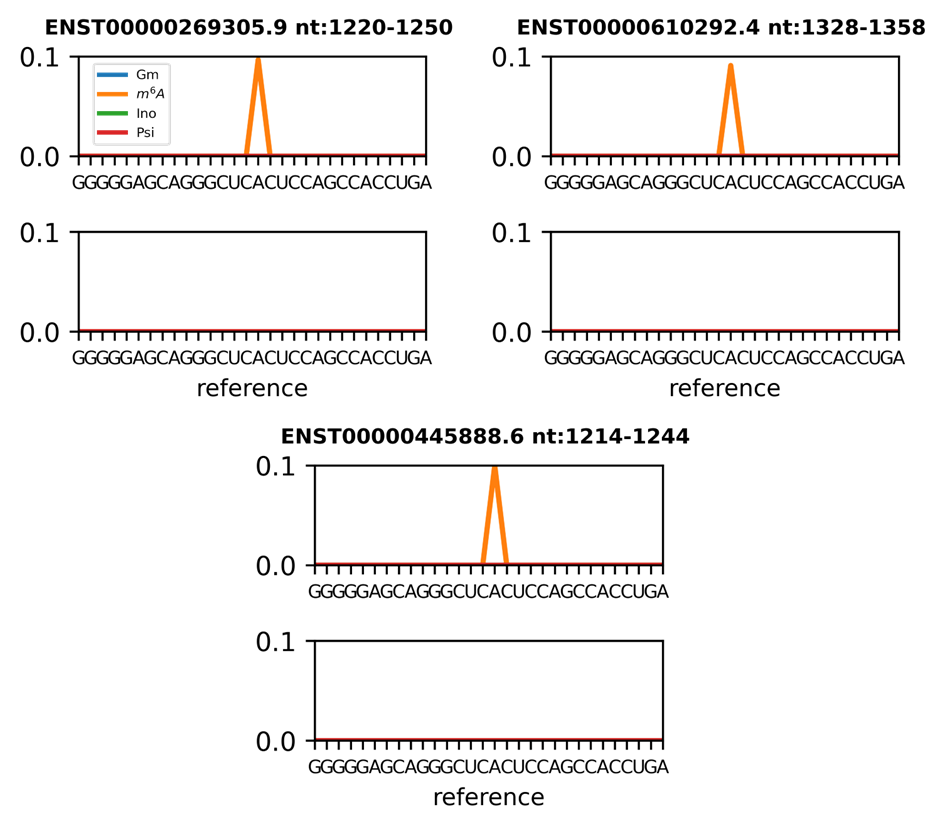

### Supplementary Fig. 8: Human blood RNA analysis using ModiDeC. Additional m^6^A sites that ModiDeC was able to detect during the analysis on the gene TP53. These peaks are currently not annotated in RMBase v3.0 database. Although further validation analyses are needed for these peak values, we found them interesting enough to show for follow-up work.

**Supplementary Fig. 9:** a–e) Examples of annotated m^6^A peaks (black dashed lines) from the atlas database that were detected by ModiDeC. f) Table showing the annotated m^6^A peaks from the atlas database detected by ModiDeC. The detected motif (the orange letter represents the m^6^A modification), chromosome, and annotated and predicted chromosome positions are shown.

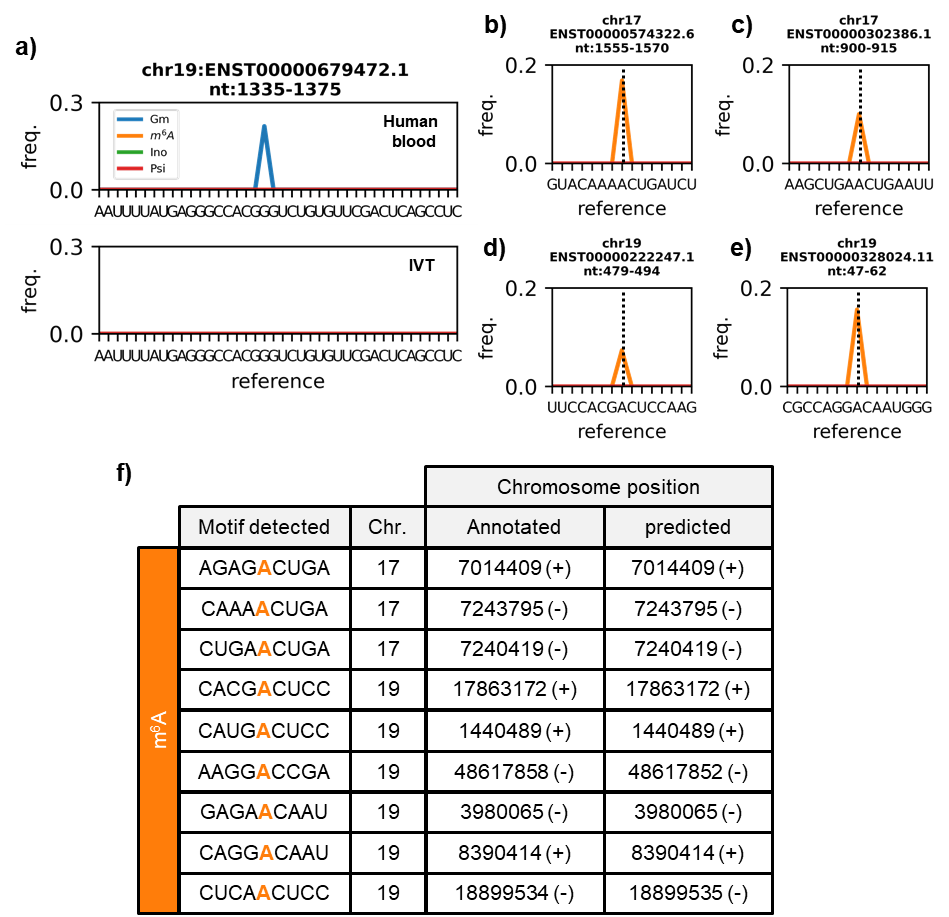

Supplementary Fig. 9 shows additional m^6^A positions identified by ModiDeC that were also confirmed by the m^6^A-Atlas database. Supplementary Fig. 9 a-d shows four of the nine positions obtained by ModiDeC with their corresponding modification frequency. Figure 9f shows a table summarizing the ModiDeC results. These are the recognized motifs, the annotated and the predicted chromosome positions. Additional analysis and the broader large transcript reference region are shown in Supplementary Fig. 10.

**Supplementary Fig. 10:** ModiDeC analysis on human blood RNA. a) ModiDeC analysis and modification frequency detection on selected transcripts with a large reference region of 1000 nt. Each subplot shows the comparison between normal and IVT human data for a specific transcript. The lowest intensity peaks in the IVT data shows that peaks in normal data are not false positives. b) 20 nt zoom of the ModiDeC analysis for a few transcripts. In this case, the x axis shows the reference nucleotides, and RNA modifications are correctly assigned to the nucleotide base to which they belong.

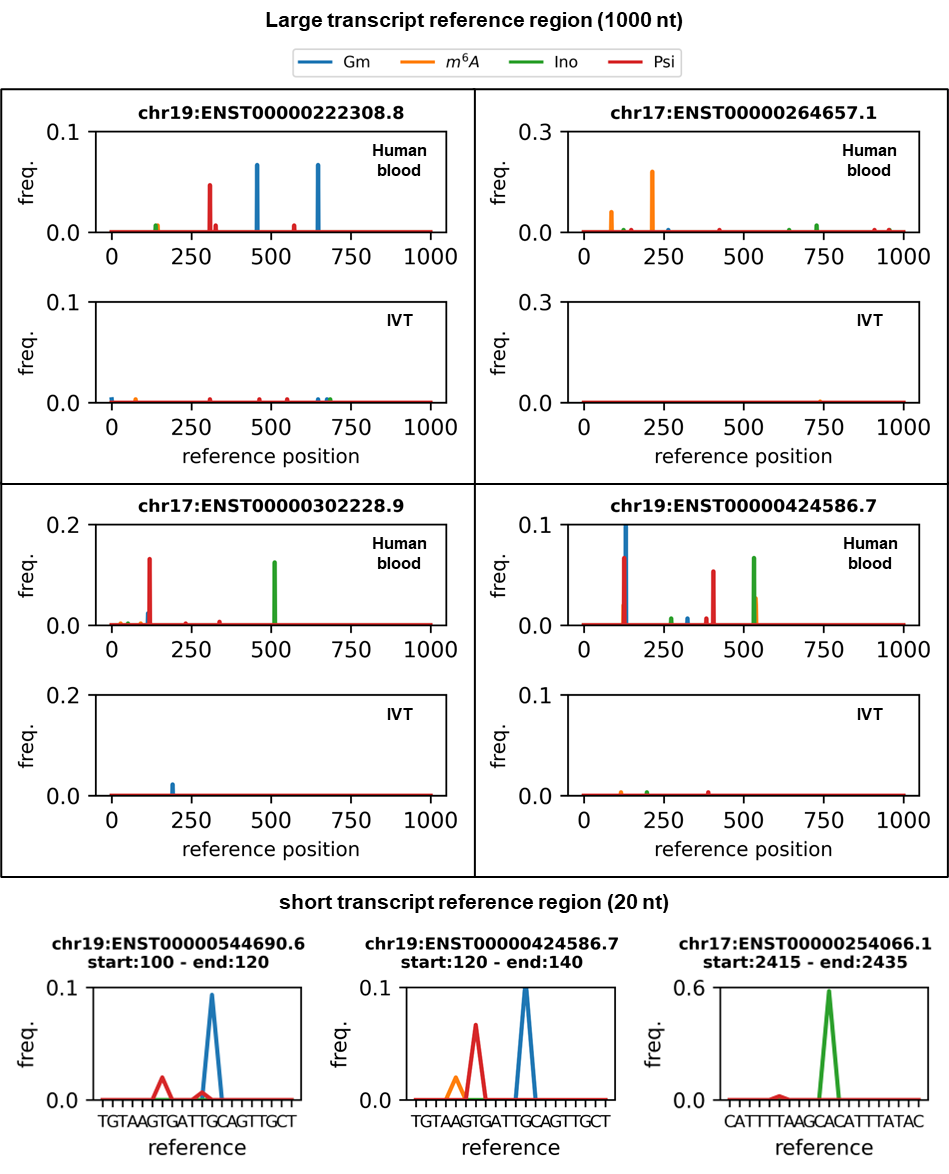

### ModiDeC RNA modification features extraction

Here we compared the transcription motifs detected by ModiDeC, that were confirmed by the m^6^A atlas database (see also Supplementary Fig. 9f), with those used for training the neural network. The comparison is shown in Supplementary Table 4. The table shows that only one of the detected motifs exactly matches a training mofit. The other detected motifs match partially or with a combination of training motifs. For example, looking the second row in the table and the motif CAAAACUGA, it can be reproduced by the CAAAA and ACUGA parts of two different training motifs. Gray letters show the missing trained nucleotides in the detected motif.

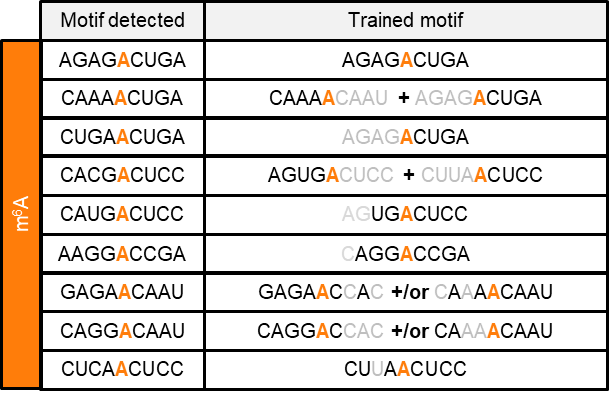

**Supplementary Table 4:** Comparison between m^6^A-modified motifs detected in ModiDeC and trained motifs. The detected motifs are not always exact matches with the trained ones. Gray letters show the missing trained nucleotides in the detected motif. The detected motifs were also confirmed by the Atlas database (see also Fig. 5g in the main text).

### SI 8. ModiDeC user graphic interface

We created three graphic user interfaces (GUIs) that can be used to generate, train, or analyze data with ModiDeC in a user-friendly environment (Fig. 6, main text). The ModiDeC GUI can be used in several ways, from retraining the neural network to directly analyzing an aligned sample using a pretrained neural network. In this section, we give an overview of how these three GUI can be used individually or all together in sequence to generate a personalized ModiDeC neural network. We decided to create the GUIs to give the opportunity to adapt and customize ModiDeC for specific problems. A full tutorial on how to use ModiDeC can be found in the GitHub page (see Code availability in the main text for the link).

Fig. 6 (main text) shows a general overview of the ModiDeC GUIs. The GUIs are titled ”ModiDeC data curation”, “ModiDeC training”, and “ModiDeC analysis”. Each section represents a different step in retraining or using ModiDeC for a final analysis.

The GUI “ModiDeC data curation” (Fig. 6, main text, top) helps the user to create data sets for training the neural network. As one can see from the graphical interface, it is possible to personalize the creation of the training data to a great extent. It is important during pre-processing to correctly assign the output labels and relative signal information for proper and correct neural network training. In fact, important features such as “modification position”, “type of modification” or “chuck size” (or sliding window size) can be personalized as input for the graphical interface for ensuring that signal-to-sequence information is stored correctly and learned during training. To start the “ModiDeC data curation” GUI, three elements are needed: 1) the pod5 folder (where measured pod5 files are stored); 2) the aligned .bam file of your data (stored in a folder); 3) the ONT level *k*-mer table for re-squiggle. Ii is important that data are base-called using the --emit-moves flag during base calling because it will change the .bam file for re-squiggle. After opening these three elements using the GUI, it is possible to set the variables that lets the user decide how to chop and label the signal during the re-squiggle. After selection, pressing the “start resquiggle” button will start the GUI generating the training data in a selected save folder. In detail, the GUI will pre-process the data in two phases. The first phase consists of the re-squiggle (signal-to-sequence assignment), where signal and the sequence are cut in personalized windows and stored in the memory. In the second phase, the modification is assigned correctly to the sequence and the output label is generated consecutively to have precisely assign a modified sequence to a signal. Next, raw signal, sequence, and labels are stored in a .npz file and they are ready to be used as training in the “ModiDeC training” section.

In the “ModiDeC training" GUI (Fig. 6, main text, middle), it is possible to retrain the neural network using the training data created in the previous step. Loading the folder of the training data using the graphical interface and setting neural networks parameters such as epochs and batch size, it is possible to start training the neural network. After the training, the trained model is saved in a selected folder and a personalized ModiDeC for a selected problem is ready to use in the analysis section.

Having a training model, it is then possible to start the “ModiDeC analysis” GUI (Fig. 6, main text, bottom). In this user interface we can load our model and pod5 and bam files to analyze the data that we are interested in. After initialization (a step to load the model and some variables), it is possible to select the number of reads to analyze and then start the neural network analysis. After pressing the button “Start analysis with neural network”, ModiDeC analyzes the raw signal of the data and results are saved in the working folder with the name “ModiDeC_analysis.npz”. To check the analysis, clicking the “visualize results" button a new window will open, and the analysis can be visualized in an interactive way.
